# Mapping SARS-CoV-2 antigenic relationships and serological responses

**DOI:** 10.1101/2022.01.28.477987

**Authors:** Samuel H. Wilks, Barbara Mühlemann, Xiaoying Shen, Sina Türeli, Eric B. LeGresley, Antonia Netzl, Miguela A. Caniza, Jesus N. Chacaltana-Huarcaya, Victor M. Corman, Xiaoju Daniell, Michael B. Datto, Fatimah S. Dawood, Thomas N. Denny, Christian Drosten, Ron A. M. Fouchier, Patricia J. Garcia, Peter J. Halfmann, Agatha Jassem, Lara M. Jeworowski, Terry C. Jones, Yoshihiro Kawaoka, Florian Krammer, Charlene McDanal, Rolando Pajon, Viviana Simon, Melissa S. Stockwell, Haili Tang, Harm van Bakel, Vic Veguilla, Richard Webby, David C. Montefiori, Derek J. Smith

**Affiliations:** Center for Pathogen Evolution, Department of Zoology, University of Cambridge, Cambridge, CB2 3EJ, UK; Institute of Virology, Charité – Universitätsmedizin Berlin, corporate member of Freie Universität Berlin, Humboldt-Universität zu Berlin, and Berlin Institute of Health, 10117 Berlin, Germany; German Centre for Infection Research (DZIF), partner site Charité, 10117 Berlin, Germany; Department of Surgery, Duke University School of Medicine, Durham, NC, USA; Duke Human Vaccine Institute, Duke University School of Medicine, Durham, NC, USA; Department of Global Pediatric Medicine, Department of Infectious Diseases, St. Jude Children’s Research Hospital, Memphis, TN, USA; Hospital Nacional Daniel A Carrión, Callao, Bellavista, Peru; Department of Pathology, Duke University School of Medicine, Durham, NC, USA; Centers for Disease Control and Prevention, Atlanta, GA, USA; Erasmus Medical Center, Rotterdam, Netherlands; School of Public Health, Universidad Peruana Cayetano Heredia, Lima, Peru; Influenza Research Institute, Department of Pathobiological Science, School of Veterinary Medicine, University of Wisconsin-Madison, Madison, WI, USA; BC Centre for Disease Control, Vancouver, British Columbia, Canada; Division of Virology, Institute of Medical Science, University of Tokyo, Tokyo, Japan; The Research Center for Global Viral Diseases, National Center for Global Health and Medicine Research Institute, Tokyo, Japan; Pandemic Preparedness, Infection and Advanced Research Center (UTOPIA), University of Tokyo, Tokyo, Japan; Department of Microbiology, Icahn School of Medicine at Mount Sinai, New York, NY, USA; Department of Pathology, Cellular and Molecular Medicine, Icahn School of Medicine at Mount Sinai, New York, NY, USA; Moderna, Inc., Cambridge, MA, USA; Division of Infectious Diseases, Department of Medicine, Icahn School of Medicine at Mount Sinai, New York, NY, USA; The Global Health and Emerging Pathogen Institute, Icahn School of Medicine at Mount Sinai, New York, NY, USA; Division of Child and Adolescent Health, Department of Pediatrics, Columbia University Vagelos College of Physicians and Surgeons, and Department of Population and Family Health, Mailman School of Public Health, New York, NY, USA; Department of Genetics and Genomic Sciences, Icahn School of Medicine at Mount Sinai, New York, NY, USA; Department of Infectious Diseases, St. Jude Children’s Research Hospital, Memphis, TN, USA

## Abstract

During the SARS-CoV-2 pandemic, multiple variants escaping pre-existing immunity emerged, causing concerns about continued protection. Here, we use antigenic cartography to analyze patterns of cross-reactivity among a panel of 21 variants and 15 groups of human sera obtained following primary infection with 10 different variants or after mRNA-1273 or mRNA-1273.351 vaccination. We find antigenic differences among pre-Omicron variants caused by substitutions at spike protein positions 417, 452, 484, and 501. Quantifying changes in response breadth over time and with additional vaccine doses, our results show the largest increase between 4 weeks and >3 months post-2nd dose. We find changes in immunodominance of different spike regions depending on the variant an individual was first exposed to, with implications for variant risk assessment and vaccine strain selection.

**One sentence summary:** Antigenic Cartography of SARS-CoV-2 variants reveals amino acid substitutions governing immune escape and immunodominance patterns.

## Main text

Since the beginning of the Severe Acute Respiratory Syndrome Coronavirus 2 (SARS-CoV-2) pandemic, the virus has caused more than 766 million cases and 6.9 million deaths (*1*). During the first year of the pandemic, circulation was dominated by the B.1 variant, characterized by the D614G substitution in the spike protein, which imparted increased infectivity and transmissibility *in vitro* and in animal models (*2, 3*) but did not escape serum neutralization (*4*). Since then, multiple variants circulated widely, with five of them B.1.1.7 (Alpha), B.1.351 (Beta), P.1 (Gamma), B.1.617.2 (Delta), and B.1.1.529 (Omicron and descendant sublineages) categorized as Variants of Concern by the World Health Organization (WHO) based on evidence of higher transmissibility, increased virulence, and/or reduced effectiveness of vaccines, therapeutics or diagnostics (*5*).

Prior to the emergence of the Omicron variants, a number of variants circulating widely in 2021 were antigenically distinct from the prototype-like 2020 viruses, with B.1.351 escaping neutralization by convalescent and post-vaccination sera the strongest (*6, 7*). Sera from individuals first infected with B.1.351 and P.1 failed to readily neutralize B.1.617.2, and vice versa, though B.1.617.2 did not show strong escape from convalescent sera after infection with prototype-like variants (*8, 9*). All variants in the Omicron lineage have substantial escape from post-vaccination and convalescent sera (*10, 11*).

It is essential to comprehend the antigenic relationships among SARS-CoV-2 variants and the substitutions that cause antigenic change. This knowledge is crucial in evaluating the need for vaccine updates and predicting whether new variants may avoid immune responses induced by current vaccines. However, with an increasing number of variants, understanding the antigenic relationships through neutralization titer data becomes more intricate. Antigenic cartography (*12*) is a tool originally developed for the analysis of human seasonal influenza virus antigenic data and has since been used in the analysis of antigenic variation in other pathogens, including avian, equine and swine influenza viruses (*13–17*), flaviviruses (*18*) including dengue viruses (*19, 20*), lyssaviruses (*21*), and foot-and-mouth disease viruses (*22*). It provides a quantitative and visual summary of antigenic differences among large numbers of variants and is a core component of the bi-annual influenza virus vaccine strain selection process convened by the WHO. Here, we use antigenic cartography to analyze patterns of cross-reactivity among a panel of 21 SARS-CoV-2 variants and 15 groups of human sera obtained from individuals following primary infection with one of ten different variants or after prototype or B.1.351 primary series vaccination. We follow this with experimental testing of point mutations to investigate the drivers of the antigenic changes observed and how the effects of subsequent changes on serological reactivity relate to the primary exposure variant.

## Results

### Variant reactivity by serum group

We used 207 serum specimens collected from vaccinated or infected individuals (table S1) and titrated in an FDA-approved (FDA/CBER Master File 026862) neutralization assay against lentiviral pseudotypes encoding the spike protein of 21 SARS-CoV-2 variants (table S2). Titrations included a panel of 15 pre-Omicron variants and six Omicron variants. Prior to the collection of each serum sample, individuals had reported no known previous SARS-CoV-2 infections or vaccination (table S1). The infecting variant was determined using whole genome sequencing. The serum specimens are from individuals with infections with D614G (n=15), B.1.1.7 (Alpha, n=14), B.1.351 (Beta, n=19), P.1 (Gamma, n=17), B.1.617.2 (Delta, n=28), B.1.526+E484K (Iota, n=6), B.1.637 (n=3), C.37 (Lambda, n=4), BA.1 (Omicron, n=7), and BA.2 (Omicron, n=1). We also included groups of sera collected from individuals twice vaccinated with a vaccine comprised of prototypic SARS-CoV-2 spike (Moderna mRNA-1273) at 4 weeks (n=32) or >3 months (n=16) post 2nd dose, and sera post 3rd dose of the same vaccine at 4 weeks (n=26) or >3 months (n=8). Finally, we included samples collected from individuals 4 weeks post 2nd dose of a B.1.351 spike vaccine (Moderna mRNA-1273.351) (n=11).

Figure 1 shows the geometric mean titer (GMT) and individual reactivity profiles for 183 sera after the exclusion of 24 outliers with titers indicative of an unreported previous infection (SOM section “Excluding outlier sera”, fig. S1). Each serum group exhibited a distinct profile of neutralization against the tested variants and, as expected, homologous serum/virus pairs were among the most potent in each group (Fig. 1A). The B.1.351 and P.1 serum groups both exhibited similar cross-neutralization of the B.1.351 and P.1 variants, suggesting a shared phenotype consistent with the closely-matched receptor binding domain (RBD) substitutions in these two variants (table S2). The Omicron variants showed the greatest escape from sera post both 2× and 3× mRNA-1273 vaccination. Notably, BA.4/BA.5 titers were substantially lower than the other Omicron variants tested in the 4-week post 3× mRNA-1273 vaccination sera, but not in the samples taken directly prior to a third dose (>3 months post 2× mRNA-1273) or >3 months post a third dose. Comparable escape was seen for the Omicron and B.1.617.2 variants against 2× mRNA-1273.351 sera. Most BA.1 sera, and the BA.2 serum sample, showed no detectable reactivity to any pre-Omicron variants. However, where titers were very high for one BA.1 serum sample, some low levels of measurable reactivity were present for the B.1.351 and D614G variants measured (Fig. 1A). Overall, our findings relating to relative antigenic escape of the different Omicron variants against different serum groups were consistent with findings in other studies that used both lentivirus pseudotype and live virus neutralization assays, where overlap was present (*11*) (fig. S6).

**Fig. 1:**
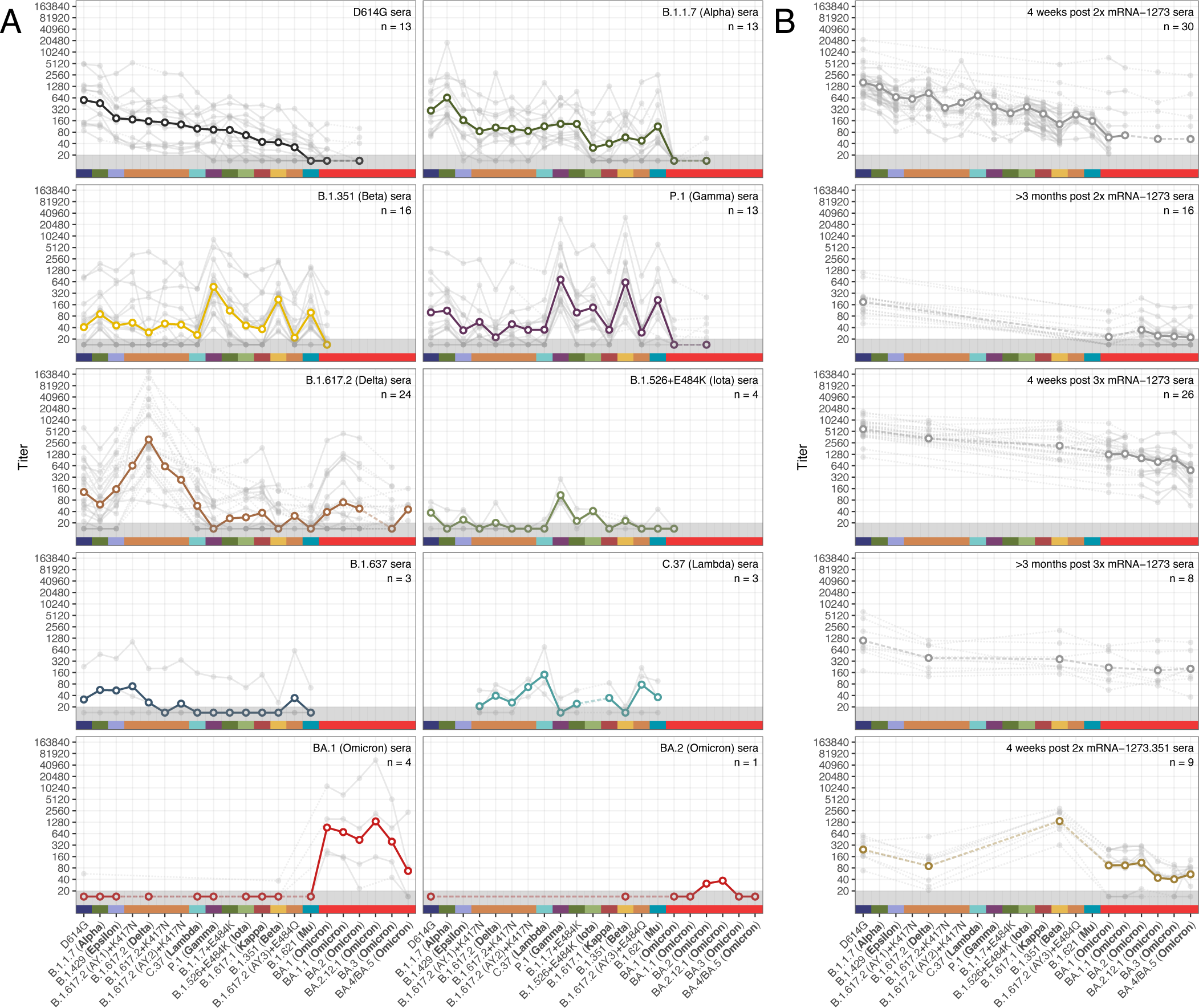
Neutralization of lentivirus pseudotypes encoding different SARS-CoV-2 spike proteins, against different groups of human sera collected after vaccination or primary infection with different variants. Serum groups are split into sera elicited by infection with different variants (**A**), and sera elicited by vaccination (**B**). Variants are ordered according to geometric mean titer (GMT) in D614G sera (panel A, top left), while additional Omicron variants are ordered chronologically. Bold lines with empty circles show the GMTs calculated after estimated differences in individual response size were removed to mitigate biases where not all sera from a group were titrated against a particular variant, as described in Materials and Methods, ‘Titer Analyses’ section. Fainter individual lines and solid points show individual serum titers. Points in the gray region at the bottom of the plots show titers and GMTs that fell below the detection threshold of 20. Each panel is labeled according to the respective serum group and color-coded as indicated on the x-axis. Fig. S2 shows titers split by sample source for the 5 serum groups where samples came from a mixture of cohorts or agencies. Titer box plots, line plots showing the individual serum titers after accounting for individual effects, and titer fold-differences relative to the homologous titer, are shown in figs. S3, S4 and S5 respectively.

Comparing post-vaccination and post-infection sera, less variation was seen in the mRNA-1273 and mRNA-1273.351 vaccine sera than in the corresponding D614G and B.1.351 convalescent sera, both in terms of response magnitude and the pattern of reactivity seen against the variants (Fig. 1, fig. S4). This is likely related to the more standardized dose (*23*) and nature of vaccination compared to infection (*24–26*) but might also be due to varying time intervals between infection and specimen collection in the convalescent specimens compared to post-vaccination specimens (*27*).

### Comparing vaccine response breadth

We calculated the breadth of post-infection and vaccination responses, controlling for differences in titer magnitude by focusing on changes in the pattern of fold-drops for each variant relative to the homologous variant (Materials and Methods, Calculating fold-drop differences in vaccine sera). We found that titer fold-drops relative to D614G in each of the mRNA-1273 vaccination serum groups had a similar pattern to D614G convalescent sera, but the size of the fold-drops was decreased by a factor, corresponding to an increased response breadth (Fig. 2A). Moreover, we found a temporal pattern of increasing response breadth (Fig. 2B), with the largest increase between 4 weeks and >3 months post 2nd dose and, to a lesser extent, between 4 weeks and >3 months post a third dose. In samples taken 4 weeks following a third vaccine dose, although titers were strongly boosted, breadth remained very similar to that measured in >3 months post 2× mRNA-1273 samples taken directly prior to the third dose. Interestingly, for the B.1.351 post-infection and vaccination groups, although titers were higher in the mRNA-1273.351 vaccination group compared to B.1.351 convalescent sera, we did not find evidence for a significant difference in the breadth of cross-reactivity (fig. S7).

**Fig. 2:**
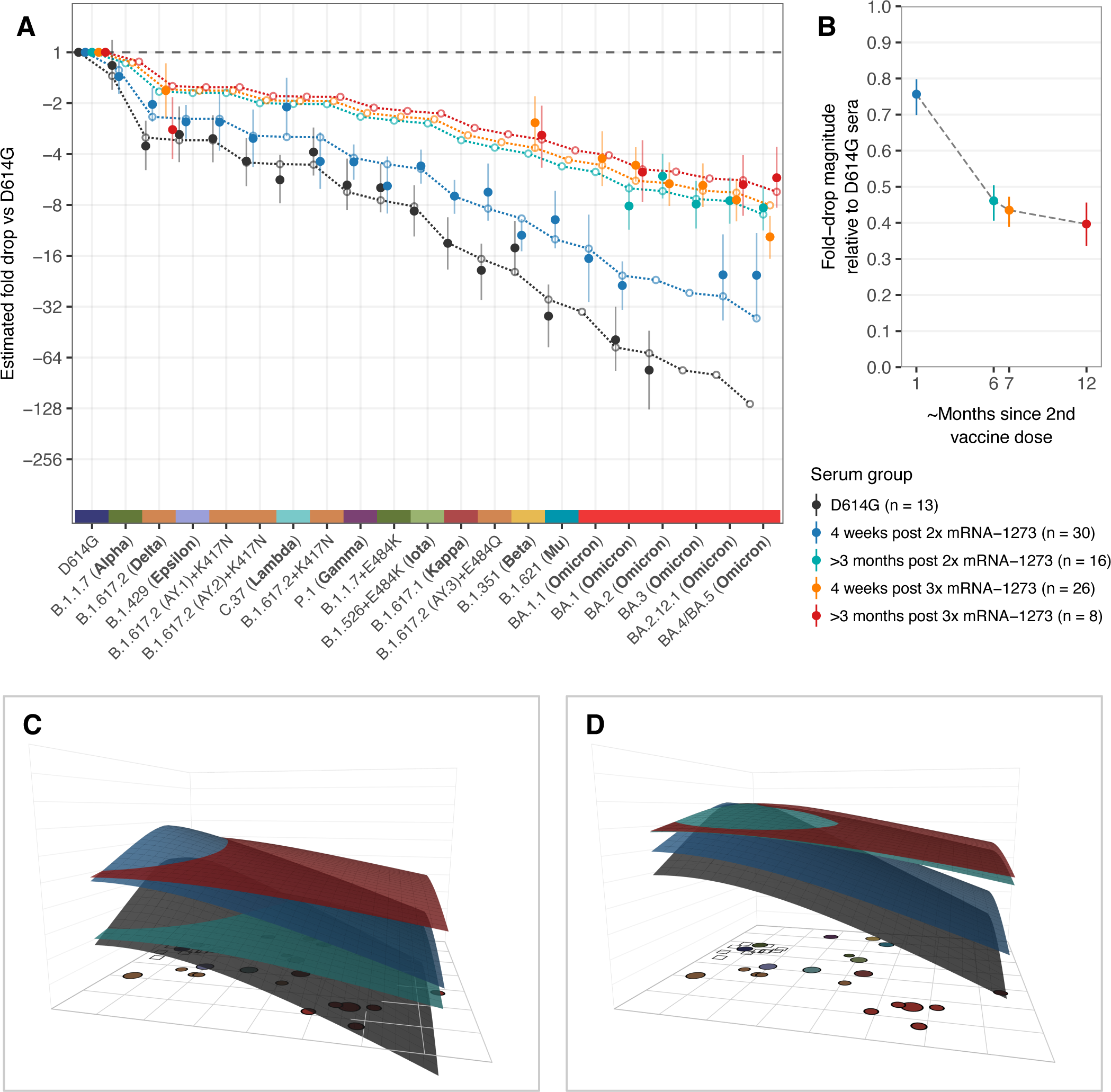
Comparison of fold-drops to different variants in post D614G infection and post mRNA-1273 vaccination sera. A) Comparison of different estimates of titer fold-drop responses against different variants. Solid points show the estimate for the mean fold drop compared to the titer for D614G, while lines represent the 95% highest density interval (HDI) for this estimate. The points for D614G to the left of the plot represents the homologous virus against which fold-change for other strains was compared and are therefore fixed at 1. Dotted lines and outline circles show estimates based on a model that assumes a shared overall pattern of fold-drops but estimates “slope” differences in the rate of reactivity drop-off seen in the 4 serum groups, as described in Materials and Methods, “Calculating fold-drop differences in vaccine sera”. To aid comparison, points and lines for each of the serum groups have some offset in the x-axis. B) Estimates of fold-drop magnitudes for each mRNA-1273 serum group, relative to the fold-drops seen in the D614G convalescent serum group. Lines show the 95% HDI for each of the estimates and the position on the x axis is proportional to the number of months since 2nd vaccine dose, assuming an average of 6 months for sera in the >3 months post 2× mRNA-1273 and >3 months post 3× mRNA-1273 groups. C) Antibody landscapes showing how estimates of the mean titer for each of the serum groups in panel A vary across antigenic space. D) Antibody landscapes as shown in C but fixed to have the same peak titer (2560 the D614G variant in order to visualize differences in the slope of the titer drop-off based on a fixed magnitude of response. Interactive versions of the landscapes shown in panels C & D are accessible online at https://acorg.github.io/mapping_SARS-CoV-2_antigenic_relationships_and_serological_responses.

### Antigenic cartography

To visualize and quantify how the different variants relate to each other antigenically, we used the titrations shown in Fig. 1 to construct an antigenic map, where antigens and sera are positioned relative to each other such that the distance between them corresponds to the fold-drop compared to the maximum serum titer (Materials and Methods, Antigenic cartography). In order to incorporate the information from the different post mRNA-1273 vaccine serum groups but also account for how their increased cross-reactivity would otherwise underestimate the relative antigenic differences between variants, we scaled distance estimates from these serum groups according to the fold-change difference estimates shown in Fig. 2B. Cross-validation results indicated that the neutralization data could generally be well represented in two dimensions (2D, fig. S9), as shown in Fig. 3A. Overall, the antigenic relationships depicted in this map were robust to assay noise and the exclusion of serum groups and variants (Materials and Methods, figs. S8-S21). The antigenic distinction between the B.1.617.2 variant and the three B.1.617.2 variants with K417N was however found to be predominantly driven by patterns of reactivity in the B.1.617.2 sera specifically (fig. S15, S16).

**Fig. 3:**
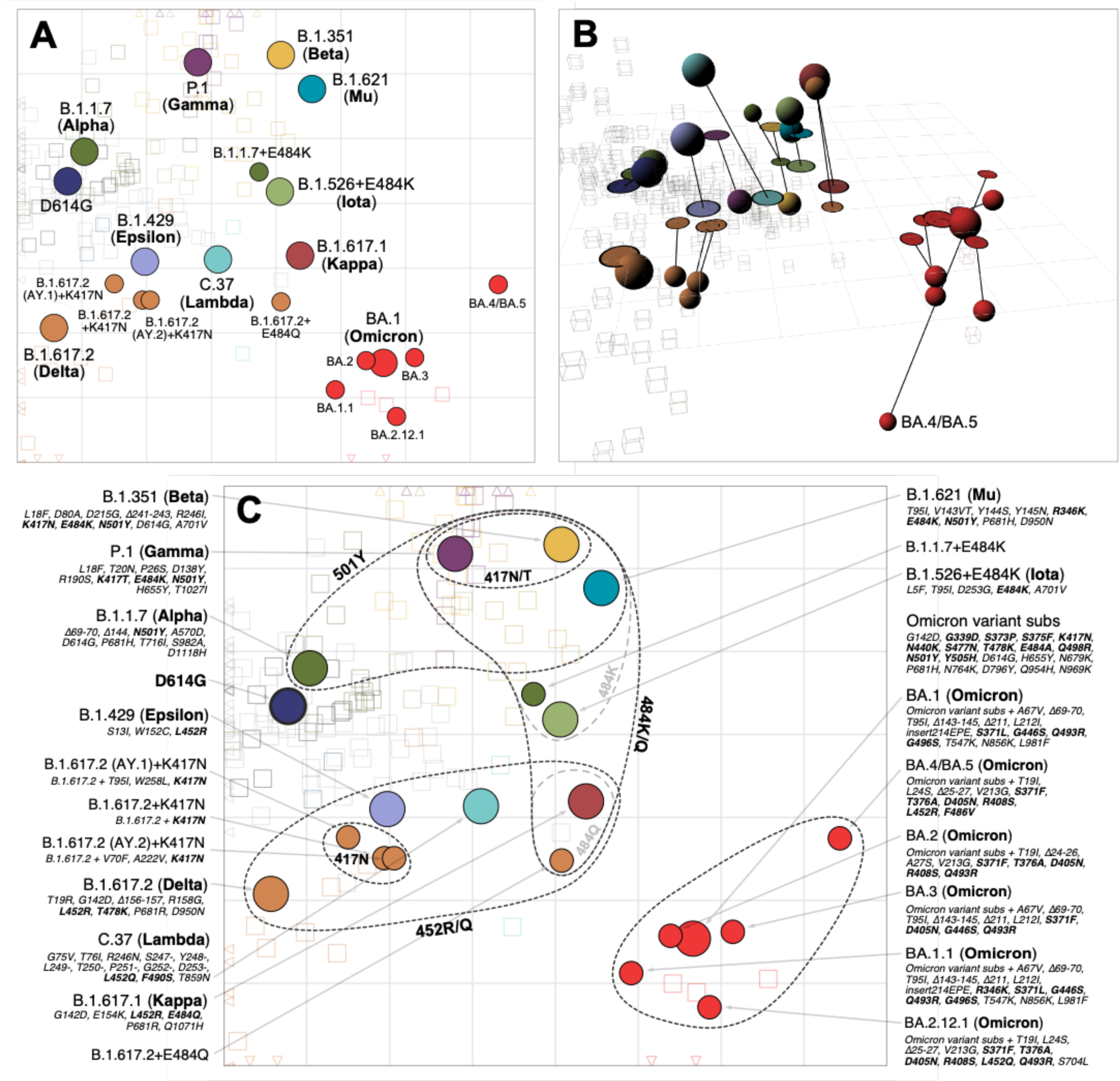
Antigenic map of SARS-CoV-2 variants and selected substitutions. A) Antigenic map with variant names. B) Antigenic map with variant positions in 3D and lines connecting to their respective positions in the 2D map. C) Antigenic map with variant names and substitutions annotated and grouped by amino acid present at spike positions 417, 452, 484 and 501, with an additional grouping for the 6 Omicron variants. Variants are shown as circles, sera as squares/cubes. Variants with additional substitutions from a root variant are denoted by smaller circles, in the color of their root variant. The x and y-axes both represent antigenic distance, with one grid square corresponding to a two-fold serum dilution in the neutralization assay. Therefore, two grid squares correspond to a four-fold dilution, three to an eight-fold dilution and so on. The x-y orientation of the map is free, as only the relative distances between variants and sera are relevant. Triangular arrowheads at the edge of the bounding box point in the direction of the sera that would be shown outside of the plot limits. A non-zoomed version of this map is shown in fig. S22. Interactive versions of the maps shown in panels A & B are available online at https://acorg.github.io/mapping_SARS-CoV-2_antigenic_relationships_and_serological_responses.

The clearest deviation from a representation of the antigenic relationships in 2D was due to the BA.4/BA.5 variant titers, for which antigenic differences fit better in 3D (Fig. 3B). Compared to 2D, the variant occupies a position that is more antigenically distinct from the other Omicron variants and closer to the B.1.617.2 sera. This positioning of BA.4/BA.5 is reflective of the fold change between BA.1 and BA.4/BA.5 in the B.1.617.2 and BA.1 serum groups (fig. S23). BA.4/BA.5 shows a substantial drop in titers compared to BA.1 in the BA.1 convalescent sera, but has increased titers compared to BA.1 in the B.1.617.2 convalescent sera, possibly because BA.4/BA.5 and B.1.617.2 share the substitutions L452R and T478K.

### Serological reactivity shown by antibody landscapes

Antigenic relationships depicted in the antigenic map provide a summary visualization of how reactivity of the different serum groups distributes amongst the variants. Figure 4 extends this visualization to antibody landscapes, where a surface in a 3rd dimension represents an estimate of how the reactivity of each serum group and individual serum varies across antigenic space (Materials and Methods, Construction of the Antibody Landscapes). The x-y plane is given by the antigenic map in Fig. 3A, while the height of the landscape over a particular point or variant represents the estimated magnitude of serum reactivity in that antigenic region. Antibody landscapes therefore give an indication of how serum reactivity distributes after exposure to different variants, how the magnitudes of the responses compare, and predicts expected levels of reactivity to variants that have not been titrated (fig. S24).

**Fig. 4:**
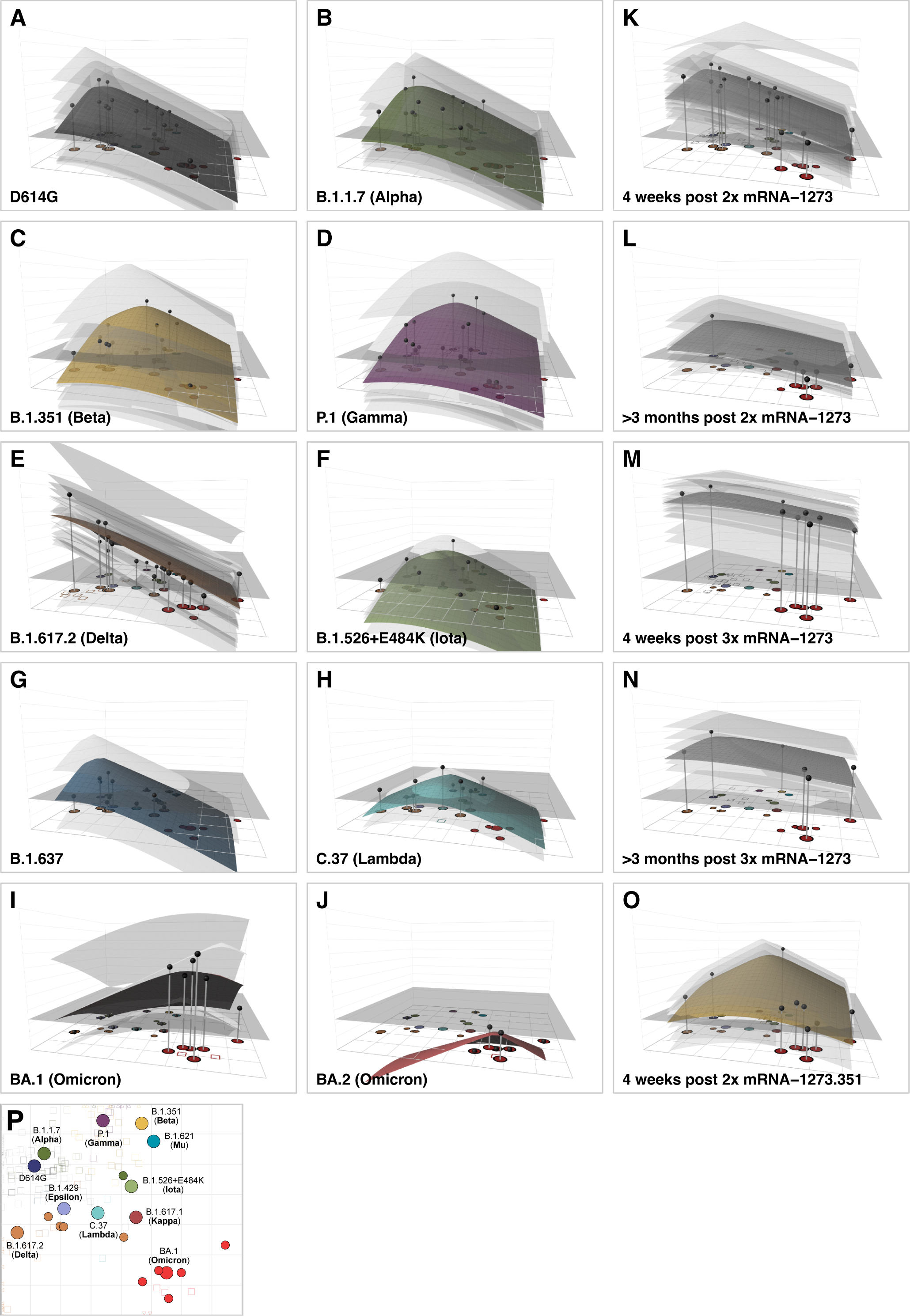
Antibody landscapes for each serum group. Colored surfaces show the GMT antibody landscapes for the different serum groups, light gray surfaces show the landscapes for individual sera. Gray impulses show the height of the GMT for a specific variant, after accounting for individual effects as described in Materials and Methods (which would otherwise bias the GMT for variants not titrated against all sera). The base x-y plane corresponds to the antigenic map shown in Fig. 3 and reproduced in panel P. The vertical z-axis in each plot corresponds to the titer on the log_2_ scale, each two-fold increment is marked, starting from a titer of 20, one unit above the map surface. The gray horizontal plane indicates the height of a titer of 50, as a reference for judging the landscapes against various estimates of neutralizing antibody correlates of protection. Additional visualizations of predicted versus fitted titers are shown in fig. S24. The number of sera included for the calculation of the landscapes are A) D614G sera (n=13), B) B.1.1.7 sera (n=13), C) B.1.351 sera (n=15), D) P.1 sera (n=13), E) B.1.617.2 sera (n=21), F) B.1.526+E484K sera (n=4), G) B.1.637 sera (n=2), H) C.37 sera (n=2), I) BA.1 sera (n=4), J) BA.2 sera (n=1), K) 4 weeks post 2× mRNA-1273 sera (n=30), L) >3 months post 2× mRNA-1273 sera (n=13), M) 4 weeks post 3× mRNA-1273 sera (n=26), N) >3 months post 3× mRNA-1273 sera (n=8), O) 4 weeks post 2× mRNA-1273.351 sera (n=9). Interactive versions of the landscapes in each of the panels are available online at https://acorg.github.io/mapping_SARS-CoV-2_antigenic_relationships_and_serological_responses.

Consistent with the titers in Fig. 1, the shape of the antibody landscapes is similar for D614G and B.1.1.7 serum groups, with the highest reactivity centered on the D614G and B.1.1.7 variants. Similarly, the shape of the landscapes generated by the mRNA-1273.351, B.1.351, and P.1 sera is comparable, with highest reactivity centered on the B.1.351 and P.1 variants. The landscape of the B.1.617.2 sera shows a contrasting topology, with the highest reactivity against the B.1.617.2 variant and falling off towards other areas of the map. On average, the B.1.617.2 sera titers against non-B.1.617.2 variants were often lower than predicted from the landscape. This suggests either that the B.1.617.2 sera could discriminate between B.1.617.2 and other variants more than was seen in reverse in the non-B.1.617.2 serum groups, or that our measurements of B.1.617.2 serum antibodies against B.1.617.2 were biased towards higher values (fig. S24). Since our data showed a larger difference in reactivity between the B.1.617.2 and D614G variants in B.1.617.2 sera when compared to other sources (fig. S6), we speculate that the latter possibility may be the case. Separately, in agreement with the titers, the landscapes show that pre-Omicron sera investigated here would be expected to have markedly reduced reactivity against variants in the Omicron lineage (Fig. 4, fig. S24). Landscapes of BA.1 first-infection sera show that the drop-off of titers to largely non-detectable levels against pre-Omicron variants is steeper than that seen in reverse for the pre-Omicron sera landscapes. This could be related to the small number of BA.1 sera or to inherent asymmetries in BA.1 and pre-Omicron sera cross-reactivity. In general, the result is consistent with other results showing that BA.1 infections generate low titers to pre-Omicron variants (*28, 29*).

The landscapes for different mRNA-1273 post-vaccination sera again illustrate how cross-reactivity differs depending on the number of, and time since, vaccinations. These differences can largely be modeled by different slopes of titer reduction across antigenic space, with reactivity that peaks in the same antigenic region but decreases at differing rates (Fig. 4K-M, Fig. 2B,C). This is true in particular for the >3 months post 2× mRNA-1273 and 4 week and >3 months post 3× mRNA-1273 serum groups for which cross-reactivity has greatly increased.

### Molecular basis of the map topology

As shown in Fig. 3C, the locations of variants in the antigenic map point to amino acid substitutions which are shared between pre-Omicron variants with similar antigenic characteristics. For example, variants with substitutions at position 484 (E to K/Q), are positioned on the right of the map due to poorer neutralization by D614G and vaccine sera. Variants towards the top of the map (B.1.1.7, B.1.351, P.1, B.1.621 (Mu), B.1.1.7+E484K) all have a substitution at position 501 (N to Y), and B.1.351 and P.1 additionally have substitutions at position 417, suggesting that these changes are associated with increased reactivity to B.1.351 and P.1 sera. Variants in the lower half of the map (B.1.617.2, B.1.429 (Epsilon), C.37, B.1.617.1 (Kappa)) all have substitutions at position 452 (L to R/Q). The Omicron variants, carrying at least 15 additional substitutions in the RBD, form a separate cluster in the lower-right of the antigenic map.

To further investigate the molecular basis of these antigenic differences, we generated 10 lentivirus pseudotypes with single substitutions at positions 417, 452, 484, and 501 in different RBD contexts and measured the effect on reactivity to different serum groups and the subsequent positioning of variants in the antigenic map. As shown in Fig. 5, in general, the antigenic effect of the different substitutions when introduced in isolation was consistent with that inferred from the antigenic map of wildtype variants. Large antigenic effects were seen for substitutions at position 484, which were associated with the right-left antigenic variation seen in the map, with D614G+E484K showing greater escape than D614G+E484Q in D614G sera (Figure 5A, fig. S25A,B). Smaller but significant effects were seen for substitutions at position 417, which was shown to mediate some of the top-bottom map variation (Figure 5B, fig. S26). Introduction of the N501Y substitution alone into D614G did not mediate large antigenic changes in the map but did cause significantly increased reactivity to B.1.1.7, P.1 and B.1.351 sera (fig. S27) and generated a virus that was antigenically similar to B.1.1.7, which is identical in the RBD (Figure 5A, table S2).

**Fig. 5:**
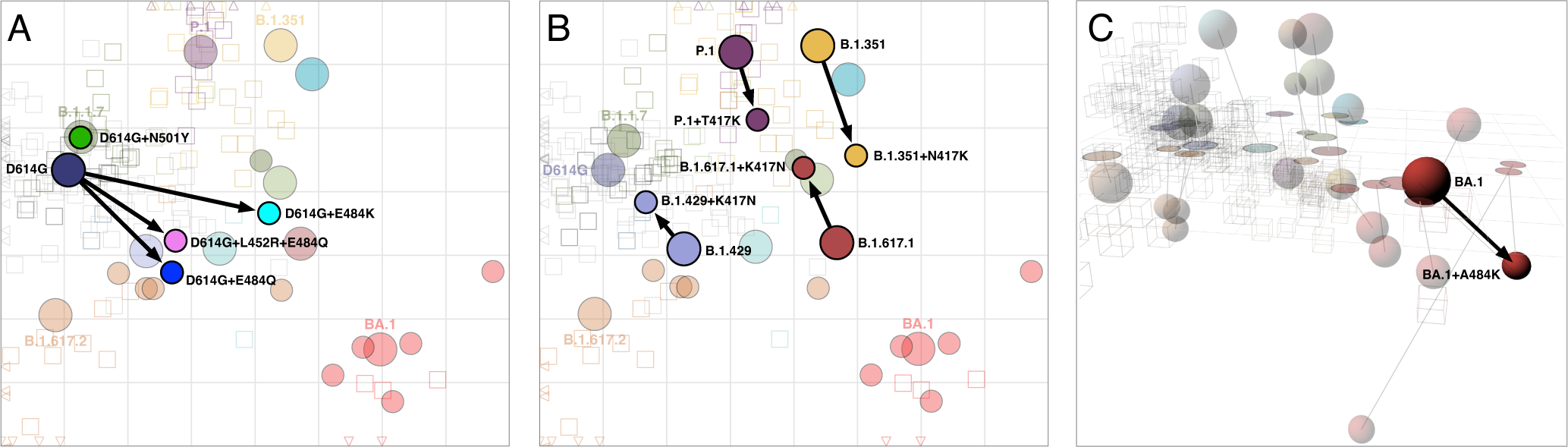
Antigenic maps including laboratory-made mutants with substitutions at positions 417, 452, 484, and 501. A) Variants with substitutions N501Y, E484K, E484Q, and L452R+E484K in the background of D614G; D614G+L452R is not shown since it was titrated against only D614G sera, so its position could not be determined. B) Variants with the T/N417K substitution in the background of P.1 and B.1.351 respectively, and K417N in the background of B.1.429 and B.1.617.1. C) BA.1 with the substitution A484K. The map in panel C is in 3D to highlight the antigenic differences between BA.1, BA.1+A484K, and BA.4/BA.5. The 2D version of panel C is shown in fig. S29. Arrows point from the antigenic position of the root virus to that of the laboratory-generated variant. Interactive versions of the maps shown in each panel are available online at https://acorg.github.io/mapping_SARS-CoV-2_antigenic_relationships_and_serological_responses.

Despite overall correspondence with map-based predictions, some results were not as expected. In particular, although D614G+L452R had significant effects decreasing reactivity to D614G sera (fig. S25C), the D614G+L452R+E484Q mutant did not show any significant difference in reactivity compared to the D614G+E484Q mutant (fig. S25B,D). Further, the B.1.429+K417N mutant showed increased reactivity to the D614G sera (fig. S26D), even though this represents a change away from sequence homology with D614G. Since it has been shown that the presence of the K417N substitution alone in the absence of 501Y greatly reduces angiotensin-converting enzyme 2 (ACE2) affinity in some contexts (*30*), we speculate that the effect seen in B.1.429+K417N may be influenced by an artificial inflation of titers generally.

Finally, the introduction of the A484K substitution into the BA.1 context allowed us to compare the effect of the alanine (A) substitution present in the Omicron variants in contrast to lysine (K) substitution seen in pre-Omicron antigenic escape variants such as B.1.351. Interestingly, we found that the BA.1+A484K substitution caused a greater escape from 4 weeks post 2× mRNA-1273 vaccine serum reactivity and also from B.1.617.2 sera (fig. S28), raising the question as to why this alternative substitution has not been seen more frequently in Omicron variants in nature. In terms of movement in the antigenic map, in 2D the BA.1+A484K mutant moves to a location to the top-right of the BA.1 variant, bringing it closer to the B.1.351 sera but also consequently close to BA.4/BA.5 (fig. S29). However, antigenic relations with BA.4/BA.5 are again better described in 3D (Fig 3B), where the BA.4/BA.5 variant utilizes the third dimension and the BA.1+A484K mutant occupies a novel area of antigenic space distinct from the other Omicron variants (Fig. 5C).

### Variation in immunodominance of different RBD sites between serum groups

Overall, a clear pattern in the substitutions tested was that not all serum groups were equally sensitive to changes at a given position, with some substitutions having a large effect on reactivity to certain serum groups but little to no effect in others. For example, although the D614G+E484K and D614G+E484Q substitutions had large effects on reactivity to D614G sera, no significant effect on B.1.351 serum reactivity was found (fig. S25A,B). Such findings are consistent with variation in immunodominance patterns and the extent to which antibodies in different sera target different structural regions in the RBD.

We tested for additional evidence of such immunodominance switches by analyzing results in the mutants alongside differences in serum reactivity between other pairs of variants that differed by single amino acid substitutions in the RBD. Figure 6A shows a summary of these comparisons for the serum groups and RBD substitutions for which the most information was available, alongside information on how pairwise differences relate to the amino acid present at that position in the serum group homologous variant. Figure S30 shows the same information across all serum groups for all single amino acid difference comparisons.

**Fig. 6:**
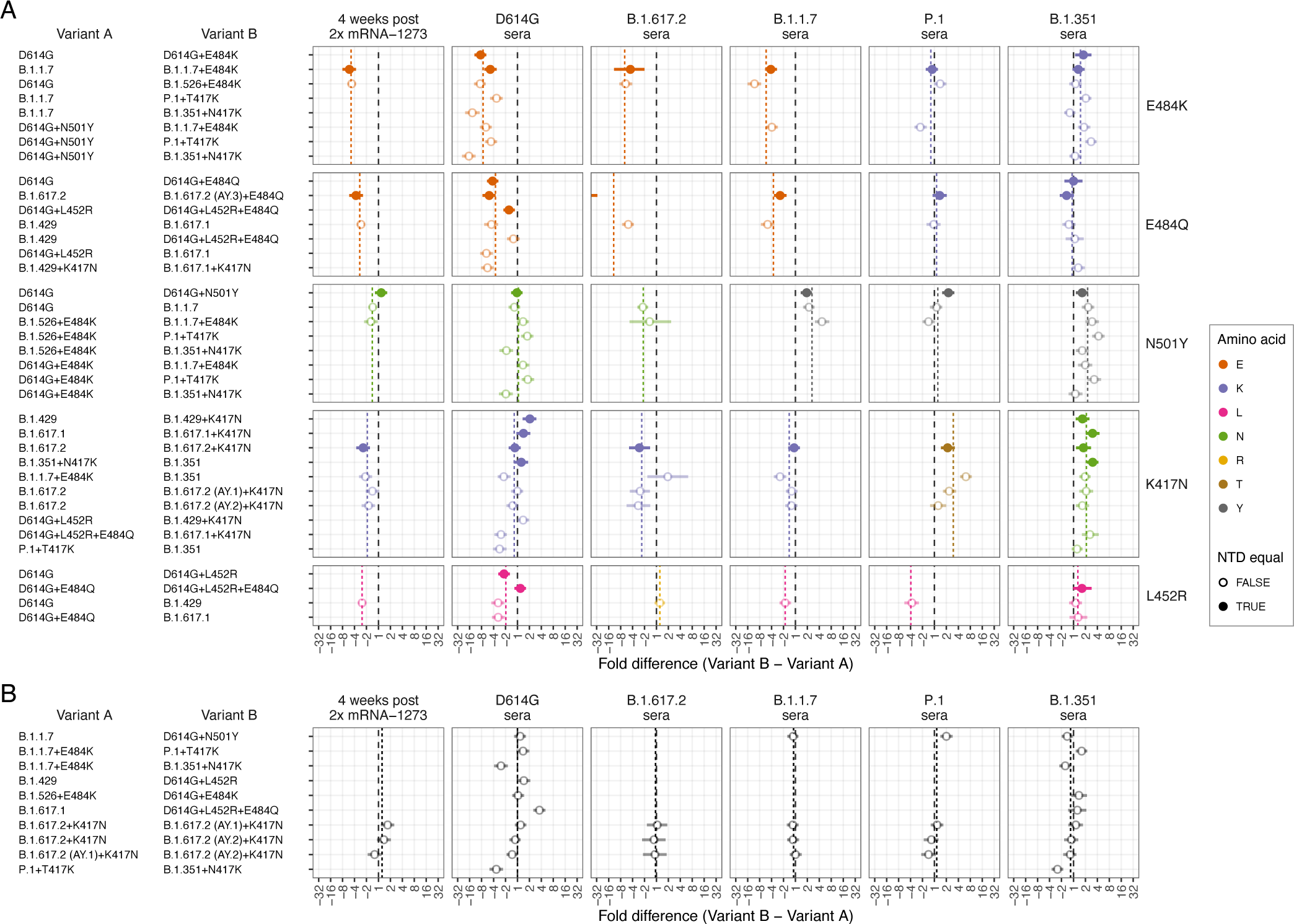
Effect of pairwise amino acid differences on reactivity to different serum groups. This plot compares the average fold difference in titer between A) different pairs of variants that differ by only a single amino acid difference in the RBD, or B) that do not differ by any amino acids in the RBD, but differ in the NTD. Comparisons are grouped by serum group (panel columns) and corresponding RBD difference (panel rows). In each panel the circle represents the estimate for the average fold difference in titer between variant A and variant B, as named on the left-hand side of the plot, while lines extend to indicate 95% highest density interval (HDI) for this estimate. The black dashed line marks a fold difference in titer of 1 (no difference), while the colored dashed line indicates the average fold difference between all pairs of variants with that substitution in the RBD. Points and lines are colored according to the amino acid in the variant homologous to that serum group, at the position in the RBD where the pair of variants compared differ. Filled circles indicate where pairs of variants had no additional amino differences in the NTD region, often because one was generated as an artificial mutant. In contrast, open circles indicate pairs of variants with additional amino acid differences in the NTD region, in addition to the RBD amino acid difference listed. The estimate for B.1.617.2 sera fold differences between the B.1.617.2 and B.1.617.2 (AY.3)+E484Q variants (panel A 3rd column, 2nd row), which falls outside the plot, is -59.4 (95% HDI -117.1, -30.7). Details of how fold-difference estimates and highest density intervals were calculated are described in Materials and Methods. Fgure S30 shows the same results against an expanded set of pairwise amino acid differences. Interactive scatterplots comparing titers against each pair of variants for each serum group are available online at https://acorg.github.io/mapping_SARS-CoV-2_antigenic_relationships_and_serological_responses.

For all pairs of variants with differences in the RBD only at position 484 from E to K or Q (Fig. 6A, rows 1 and 2), D614G sera consistently showed significantly reduced reactivity, while the B.1.351 sera showed little to no significant change in reactivity. Across the other serum groups, a general pattern was that sera from individuals infected with variants with the ancestral 484E (2x mRNA-1273, D614G, B.1.1.7, and B.1.617.2) were sensitive to differences at position 484, while sera from infections with variants with 484K (B.1.351, P.1, and B.1.526+E484K) were more resistant to these substitutions, although B.1.351 did show evidence for some smaller increases in titers in some cases. The BA.1+A484K mutant also reflected this pattern of sensitivities in the serum groups against which it was titrated, with significant titer decreases in mRNA-1273 and B.1.617.2 sera (484E) and a smaller increase in reactivity to mRNA-1273.351 sera (484K) (fig. S28, S30). Of the four BA.1 sera (484A), the mean fold-decrease of 2.1 against the BA.1+A484K mutant versus the BA.1 variant suggests that these sera are also sensitive to changes at the 484 position.

Where variants differed only by N501Y (Fig. 6A, row 3), we also found that sensitivity was linked to the amino acid at position 501 in the infecting variant. The two serum groups resulting from infections with B.1.351 and B.1.1.7 (both with 501Y), were sensitive to differences at position 501, while the serum groups post-Prototype vaccination and post-D614G, B.1.526+E484K, and B.1.617.2 infection (variants with ancestral 501N), were typically not sensitive to the N501Y substitution.

For K417N comparisons (Fig. 6A, row 4), there was evidence for equal or decreased titers in serum groups with exposure to variants with the ancestral K at position 417 (mRNA-1273, D614G, B.1.617.2 and B.1.1.7 serum groups), and a corresponding overall increase in titers to the serum group post infection with B.1.351 (417N). P.1 sera (417T) showed increased reactivity to the K417N substitution. The B.1.351 sera (417N) also showed increased reactivity associated with the K417T substitution, likely reflecting a structural homology of the changes caused by K417T and K417N.

Finally, where variants allowed for a comparison of the effect of the substitution L452R (Fig. 6A, row 5), we found evidence for most of the serum groups distinguishing between variants that differed by this substitution, with decreased (mRNA-1273, D614G, B.1.1.7, and P.1 sera) or increased (B.1.617.2 sera) titers, corresponding to the amino acid present at position 452 in the eliciting variant. One exception was the B.1.351 sera, where, unlike the other serum groups with the ancestral 452L, the L452R substitution did not produce an overall decrease in titers.

### Impact of changes in the NTD

We also investigated the effects of substitutions in the N-terminal domain (NTD). Generally, the effects of single amino acid differences in the RBD were consistent regardless of whether additional NTD differences were present or not (Fig. 6A). This was also reflected in pairwise comparisons of variants with no RBD differences, where we typically found no significant differences in titer reactivity (Fig. 6B). This included little evidence for a significant effect on serum reactivity of features such as the NTD 69-70 deletion in the B.1.1.7 variant (Fig. 6B, B.1.1.7 vs D614G+N501Y). However, for comparators involving the B.1.351+N417K mutant, which has the substantial B.1.351 NTD changes that include the 241-243 deletion and R246I substitution, we did find evidence of differences to other viruses that had sequence homology in the RBD such as the P.1+T417K and B.1.1.7+E484K mutants. Although these differences may be associated with effects of these NTD substitutions on titer reactivity (as has been found in other cases (*31*)), we found that in the comparisons shown in Fig. 6B, the B.1.351+N417K mutant also had lower titers against the B.1.351 sera themselves, despite sequence homology of the NTD differences for these sera. This may suggest that in our data there was a general negative bias in titers measured against the B.1.351+N417K mutant, rather than antigenic effects of the substitutions within the NTD.

## Discussion

The antigenic analyses of SARS-CoV-2 variants presented here underscore the advantages of an integrated and extensible framework for understanding antigenic relationships and serum responses in SARS-CoV-2. The antigenic map and the antibody landscapes allow the comparison of serum responses not just on a per-variant basis, but also provide a quantitative measurement of both the magnitude and breadth of the response following different exposures, including against variants that have not been measured. Although the sera investigated here represent exposures to a single variant, the same principles can be applied for understanding and comparing how multi-exposure serum responses distribute across antigenic space (*32*), relevant for ongoing studies seeking to compare the immunity built through different prospective vaccination regimens (*33, 34*).

Using regression analyses and antibody landscapes, we introduce a method to quantify and visualize changes in response cross-reactivity breadth, disentangling it from changes in cross-reactivity due to differences in response magnitude. Applying this approach to mRNA-1273 vaccination samples taken at multiple time points allows us to quantify how response breadth changes over time, independent of boosting and waning of raw titer magnitude (Fig. 2). In particular, the similar estimates of response breadth just prior to the 3rd vaccination, and from 4 weeks post 3rd vaccination, show that the main short-term effect of the 3rd vaccination was to boost the magnitude of a response that had already become more cross-reactive, rather than to generate significant additional breadth of cross-reactivity. Notably, this is consistent with studies into short-term influenza vaccine responses, where the predominant effect of vaccination is boosting of pre-existing patterns of pre-vaccination antibody reactivity (*32*). In influenza, this boosting effect is independent of the vaccine variant used. It is possible that the third vaccine dose was responsible for the later additional increases in breadth in >3 months post-3rd dose samples, though the effect size was small and not statistically significant. It is also possible that the process we observe of increasing response breadth over time (Fig. 2B) would have continued to this time point without a third vaccination. This being said, increases in breadth in the absence of a third dose would not have translated to increased protection without the effect of the 3rd dose boosting titer magnitude.

We also find that different variants and serum responses cluster on the antigenic map in a way that allows the inference of prime candidates for the amino acid substitutions responsible for the antigenic changes. The significant impact of substitutions at position 484 and the role of 417 and 501 agree with other work based on deep mutational scanning (*35–37*) and neutralization data using vesicular stomatitis virus pseudotyped particles (*38, 39*). The large antigenic effect of single substitutions such as E484K is reminiscent of human, swine, and equine influenza viruses, where, among circulating strains that may differ by amino acids at multiple positions, a large portion of the antigenic difference is associated with only a single or double substitution (*40–42*). For influenza virus, these substitutions are located adjacent to the receptor binding site on the hemagglutinin protein, which is also observed for the determinants of SARS-CoV-2 antigenic evolution described here, being located close to the ACE2 binding site on the spike protein (fig. S31, S32). Additionally, of the 12 RBD substitutions shared between the Omicron variants tested, S371F/L, N440K, E484A, and Q493R have been found to have significant effects on monoclonal antibody neutralization (*36, 39, 43*), in addition to the substitutions K417N and N501Y, which are already implicated as relevant for inhibiting neutralization in other variants (*37, 44, 45*). Except for S371F/L and N440K, these substitutions also follow a pattern of ACE2 binding site proximity. The broad effects of S371L on monoclonal antibodies binding to multiple epitopes on the spike suggests there may be structural cascade effects of that particular substitution that also affects regions closer to the ACE2 binding site (*39*).

Our observations relating to immunodominance changes of different antigenic regions have been seen similarly in the influenza virus (*46*), and are consistent with studies in SARS-CoV-2 showing that monoclonal antibodies tend to have different regional binding preferences when sourced from individuals exposed to different variants (*9, 47–49*). Conclusions of an immunodominance switch for B.1.351 sera away from the 484 region also correspond well with published data applying deep mutational scanning techniques to sera post D614G and B.1.351 infection (*37*). Here, our results show how such immunodominance switches extend to vaccination and infections with different variants and how changes can be associated with the amino acid present in the different eliciting variants. These observations explain the wider patterns in the data where certain serum groups may distinguish clear antigenic differences between certain variants while others may not, and again underscore that antigenic differences between variants are not necessarily absolute measures, but can be relative to the particular sera against which variants were titrated and the specific structural regions that the particular sera tested predominantly target.

We note two limitations of the analyses presented here. First, although the assay we use is FDA-certified, it measures neutralization using lentivirus pseudotypes, which may differ from live virus neutralization assays. In this regard, where there was overlap with other reported data, we generally find our fold-difference estimates to be within the reported ranges. In particular, we did not find an obvious bias according to comparisons with live-virus assay results (fig. S6), in keeping with other comparisons (*11*). Second, we focus here on patterns of serological reactivity and antigenic variation seen only up until the early Omicrons. Beyond this, it is very difficult to source sufficient primary exposure human sera to study antigenic relationships in detail, and it is increasingly necessary to rely on animal model sera. Consequently, it is critical to assess whether the patterns of antigenic relatedness and changes in immunodominance found for human sera can be reliably measured in different animal models.

As populations increasingly experience multiple exposures, it will also be important to investigate how antigenic differences according to multi-exposure serum responses compare with those inferred from primary exposure sera, in particular with immunodominance patterns in mind. For example, findings that infection with Omicron BA.1 after a previous exposure to pre-Omicron variants predominately boosts antibodies targeting epitopes shared between pre-Omicron and Omicron variants highlight the significance of understanding how different initial exposures dictate the structural regions that were initially targeted, and how this interacts with a future exposure (*50–53*). Answers to these questions will better reveal the variants and substitutions to which different populations are most vulnerable and help anticipate which emerging variants may be most at risk of evading current immunity. The choice of vaccine immunogens based on immunodominance considerations may be as important as their antigenic characteristics.

## Funding

SHW, ST, EBL, AN, RAMF, TCJ, DJS: NIH Centers of Excellence for Influenza Research and Surveillance (CEIRS, contract #HHSN272201400008C) and Centers of Excellence for Influenza Research and Response (CEIRR, contract #75N93021C00014).

XS, DCM, CM, HT, LD: SARS-CoV-2 Assessment of Viral Evolution (SAVE) Program (Contract #75N93019C00050 Opt 18A & 18B).

BM, TCJ, CD, VMC: Funded by the German Federal Ministry of Education and Research through project DZIF (8040701710 and 8064701703) and the German Federal Ministry of Health through project SeroVarCoV.

VMC, LMJ: Funded by the German Federal Ministry of Education and Research through project VARIpath (01KI2021)

HvB, VS, FK: NIAID Centers of Excellence for Influenza Research and Response (CEIRR) contract 75N93021C00014, SAVE option 12A, the NIAID Collaborative Influenza Vaccine Innovation Centers (CIVIC) contract 75N93019C00051.

EBL is funded by a Gates Cambridge Scholarship.

RP: Moderna, Inc. Also supported by the Office of the Assistant Secretary for Preparedness and Response, Biomedical Advanced Research and Development Authority (contract 75A50120C00034) and by the National Institute of Allergy and Infectious Diseases (NIAID).

YK: Center for Research on Influenza Pathogenesis and Transmission (CRIPT) (75N93021C00014), and the Japan Program for Infectious Diseases Research and Infrastructure (JP21wm0125002) from the Japan Agency for Medical Research and Development (AMED).

MSS: Funded by the US Centers for Disease Control and Prevention (contract 75D30120C08150 with Abt Associates).

## Author contributions

Conceptualization: SHW, BM, XS, DCM, DJS

Methodology: SHW, DJS

Software: SHW, TCJ

Validation: SHW, BM, XS, ST, EBL, AN, DCM, DJS

Formal analysis: SHW, BM, XS, ST, EBL, AN

Investigation: SHW, BM, XS, ST, EBL, AN, DCM, DJS

Resources: XS, MAC, JNC, MBD, TND, CD, RAMF, PJG, VMC, LJ, PJH, AJ, YK, FK, RP, VS, MS, FSD, VV, HvB, RW, DCM, DJS, LMJ, VMC, CD

Data curation: SHW, BM, XS

Writing - Original Draft: SHW, BM, DJS

Writing - Review and Editing: All

Visualization: SHW, BM, ST, AN, EBL

Supervision: DCM, DJS

## Disclaimers

The findings and conclusions in this report are those of the author(s) and do not necessarily represent the official position of the [US] Centers for Disease Control and Prevention (CDC).

## Data and materials availability

Code and data can be accessed from GitHub webpage: https://github.com/acorg/mapping_SARS-CoV-2_antigenic_relationships_and_serological_responses

