## Supporting Online Material for "Mapping SARS-CoV-2 antigenic relationships and serological responses"

**This PDF file includes**

Materials and Methods

Fig S1-S32

Table S1-S2

**Materials and Methods**

***Specimens and study cohorts***

All samples, including those from vaccine recipients were collected in clinical studies with information listed in Table S1. Convalescent sera are assumed but not confirmed first infections, based on no previously recorded or self-reported SARS-CoV-2 vaccination or infection, plus additional criteria in some cases as also described in Table S1. For convalescent samples, the infecting variant was determined using whole genome sequencing. Institutional review board (IRB) approvals were obtained for each study site. All participants were given and signed the informed consent form.

Vaccination samples came from a variety of sources, as also indicated in Table S1 and described in more detail below:

*4-weeks post 2x mRNA-1273*

The majority (n=28) of the 4-week post 2x mRNA-1273 samples were from the mRNA-1273 phase 1 study (NCT04283461) and were obtained from the Division of Microbiology and Infectious Diseases (DMID), National Institute of Allergy and Infectious Diseases (NIAID) for the mRNA-1273 phase 1 study team and Moderna Inc. The phase 1 study protocols and results are reported previously (*54*, *55*). The phase 1 trial tested the identical vaccine (mRNA-1273), dose (100 µg) and schedule as used in the Moderna phase 3 (NCT04470427); titrations against these samples were also reported in (*56*). A smaller subset of samples (n=4) were obtained from the Coronavirus Efficacy (COVE) phase 3 trial, and were pre-selected as having high titers of neutralizing antibodies against D614G in order to enable detection of neutralization against the wider range of Omicron variants against which they were measured; some titrations against these samples also appear in (*57*). Finally, an additional 11 sera from the COVE phase 3 trial were titrated against D614G and the D614G+N501Y mutant, and an additional 26 sera were titrated against BA.1 and the BA.1+A484K mutant. These sera appear in the results shown in fig. S27 and S28, respectively, but are not included in the general analysis due to lack of titrations against further variants.

*>3-months post 2x mRNA-1273*

All samples relate to sera previously described in (*58*) that were obtained before homologous mRNA-1273 boosting (50 µg) in 16 study participants who had received two inoculations of mRNA-1273 (100 µg) under emergency-use authorization. These participants were part of a subset analysis of high D614G responders in the NIAID heterologous boost study DMID 21-0012.

*4 weeks post 3x mRNA-1273*

16 samples relate to sera from the same individuals in the >3-months post 2x mRNA-1273 group, but obtained 29 days after homologous mRNA-1273 boosting (50 µg), again as described and appearing in (*58*). An additional 10 samples also relate to samples collected as part of the DMID 21-0012 and also relate to samples previously described in (*57*), which were also a subset of high D614G responders.

*>3 months post 3x mRNA-1273*

These samples were taken as part of the COVID-19 Variant Immunologic Landscape (COVAIL) randomized clinical trial (NCT 05289037) and are from the subset of individuals who had received a previous mRNA-1273 vaccine regimen followed by a mRNA-1273 booster dose. Samples were taken directly prior to a 4th mRNA-1273 dose and, as for other vaccine groups, we subset to those with no previous record of a SARS-CoV-2 infection.

*4 weeks post 2x mRNA-1273.351*

Samples were from COVID-19 naïve participants who received two 100 µg doses of mRNA-1273.351 on days 1 and 29 as part of a clinical trial (NCT 04785144), experimental arm 2C.

The use of the specimens for the research conducted in this manuscript is covered under IRB protocols Pro00093087 and Pro00105358.

Where samples came from the same serum group came from a mixture of cohorts or agencies, we did not find evidence for differences in the patterns of reactivity according to cohort or agency (fig. S2).

***Pseudotype virus assay***

Neutralization was measured with lentiviral particles pseudotyped with SARS-CoV-2 spike and containing a firefly luciferase (Luc) reporter gene for quantitative measurements of infection by relative luminescence units (RLU). The assay was performed in 293T/ACE2-MF provided by Drs. Michael Farzan and Huihui Mu. Pseudoviruses were prepared, titrated, and used for measurements of neutralizing antibodies essentially as described previously (*56*). Briefly, an expression plasmid encoding codon-optimized full-length spike of the Wuhan-1 ancestral sequence (VRC7480) was provided by Drs. Barney Graham and Kizzmekia Corbett at the Vaccine Research Center, National Institutes of Health (USA). Mutations were introduced into VRC7480 either by site-directed mutagenesis using the QuikChange Lightning Site-Directed Mutagenesis Kit from Agilent Technologies (Catalog # 210518), or were created by a spike gene synthesized by GenScript using the spike sequence in VRC7480 as template. All mutations were confirmed by full-length spike gene sequencing by Sanger Sequencing, using Sequencher and

SnapGene for sequence analyses. Pseudovirions were produced in HEK293T/17 cells (ATCC cat. no. CRL-11268) by transfection using Fugene 6 (Promega Cat#E2692) and a combination of spike plasmid, lentiviral backbone plasmid (pCMV ΔR8.2) and firefly Luc reporter gene plasmid (pHR' CMV Luc)36 in a 1:17:17 ratio in Opti-MEM (Life Technologies). Transfection mixtures were added to pre-seeded HEK 293T/17 cells in T-75 flasks containing 12 ml of growth medium and incubated for 16-20 h at 37°C. Medium was removed and 15 ml of fresh growth medium added. Pseudovirus-containing culture medium was collected after an additional 2 days of incubation and clarified of cells by low-speed centrifugation and 0.45 μm micron filtration. TCID50 assays were performed as described previously (*56*).

For measurements of neutralization, a pre-titrated dose of virus was incubated with 8 serial 5-fold dilutions of serum samples (1:10 or 1:20 starting dilution) in duplicate in a total volume of 150 μl for 1 h at 37°C in 96-well flat-bottom culture plates. 293T/ACE2-MF cells were detached from T75 culture flasks using TrypLE Select Enzyme solution, suspended in growth medium (100,000 cells/ml) and immediately added to all wells (10,000 cells in 100 μL of growth medium per well). One set of 8 wells received cells and virus (virus control) and another set of 8 wells received cells only (background control). After 66-72 h of incubation, medium was removed by gentle aspiration and 30 μl of Promega 1X lysis buffer was added to all wells. After a 10-minute incubation at room temperature, 100 μl of Bright-Glo luciferase reagent was added to all wells. After 1-2 minutes, 110 μl of the cell lysate was transferred to a black/white plate. Luminescence was measured using a GloMax Navigator luminometer (Promega). Serum samples were heat-inactivated for 30 minutes at 56ºC prior to assay. Neutralization titers are the inhibitory dilution (ID) of serum samples at which RLUs were reduced by either 50% (ID50) or 80% (ID80) compared to virus control wells after subtraction of background RLUs. The spike mutations of all variants used in this study are shown in Table S2.

***Titer analyses***

*Excluding outlier sera*

The serum samples used in this study were collected from individuals with no reported previous vaccination or infection preceding the exposure that was associated with their respective serum group, with further additional criteria for some sample sources as described in table S1. However, as with most studies it is possible that for some individuals, additional previous (even asymptomatic) infections were not detected or reported. Based on this possibility, we sought to identify sera with patterns of reactivity that were less likely to be consistent with a true first exposure. As such, we applied selection criteria to all the sera to identify cases where a titer was measured that was higher than the serum-group-homologous-variant titer by more than 2-fold (for example a serum from the

B.1.617.2 group that had a titer against D614G that was more than 2-fold higher than the titer measured against B.1.617.2). Sub-variants of B.1.617.2 and BA.1 (identified with smaller circles in Fig.3) were excluded from this analysis, as well as the variant P.1, which we noted tended to often have high titers generally. For example, P.1 often has higher than homologous titers for sera from the B.1.351 serum group, possibly due to assay-related biases inflating titers for the particular P.1 variant, as it has been observed in other assays using lentiviral pseudotypes (*8*). B.1.429 was used as the homologous variant for sera from the B.1.637 group, being the genetically most similar variant titrated, while D614G was used for mRNA-1273 vaccination groups, and B.1.351 was used for the mRNA-1273.351 vaccination group. 15 of the total of 207 sera had titers exceeding the 2-fold-higher-than-homologous criteria described above, and are highlighted in red in the titer plot shown in fig. S1.

As an additional step we also excluded nine sera as likely second infection responses based on visual inspection of patterns of reactivity that deviated strongly from the general trend of titer reactivity for a given serum group. These sera are additionally highlighted in fig. S1 in blue. In particular, this criteria applied to sera from the BA.1 serum group where we noted a clearly bimodal pattern of reactivity with three post-BA.1 infection sera that showed high titers against the D614G variant and four sera which did not. Based on the large antigenic difference between D614G and BA.1 and the types of reactivity seen in more controlled animal model BA.1 infections (*59*), we judged the four more BA.1-specific sera to be likely first infections and excluded the rest of the group as likely cases that had had a previous exposure to a pre-Omicron variant.

*Calculation of Geometric Mean Titers (GMT) and fold-changes for non-detectable titers*

Calculation of the mean titers becomes non-trivial when experiments contain non-detectable titers or, in data science terminology, censored data. Although computation of means of data with censored values has been around for over 60 years (*60–62*), it does not seem to be commonly employed when analyzing data from neutralization assays. Correct estimation of the GMT becomes especially important if one wants to quantify the amount of escape by variants like the Omicron variants, with many non-detectable titers. Setting non-detectable titers to half the value of an assay's limit of detection is a common method employed in virology, however, it may overestimate the GMTs for such variants. Below we give a brief overview of our approach to dealing with such censored data, functions for which are included in the associated "titertools" package (*63*).

The main step in estimation of the titers is the construction of a likelihood function that describes the probability of observing both the non-detectable and detectable titers. In the dataset used for this study we deal with only left-censored data. We assume that $\log_2$ titer values of a particular serum group are modeled by a normal density function $f(x; \mu, \sigma)$ with unknown mean $\mu$ and standard deviation $\sigma$ and we let $y_i (i = 1, \dots, M)$ denote detectable titers and $\mathrm{T_i} (\mathrm{i} = 1, \dots, \mathrm{N})$ denote the non-detectable titers. Then the log likelihood function is given by

$$LL(\mu, \sigma) = \sum_{i=1}^{N} \log \int_{-\infty}^{T_i} f(x; \mu, \sigma) + \sum_{i=1}^{M} \log f(y_i; \mu, \sigma)$$

In this expression, the term involving the integral is the likelihood of observing the censored titers, whereas the other term is the likelihood of observing the non-censored titers. This likelihood function then can either be used in a maximum likelihood estimation pipeline or, if priors are chosen for $\mu$ and $\sigma$, in a Bayesian posterior calculation pipeline from which the value of $\mu$ can be estimated in various ways. For the calculations in this paper, confidence intervals were calculated using the highest density interval (HDI) method of calculation from 10000 samples.

When calculating the fold-changes in titers, the same approach to non-detectable titers was taken, with for example a change from 40 to <20 equating to a change of <-1 on the $\log_2$ scale, for which the likelihood function would be the integral of the normal density function from -∞ to -1. For estimating fold-changes we used the function “log2diff” in the “titertools” package, with parameter “sigma = 0.87”, based on our estimates of the magnitude of variation in paired measurement repeats (fig. S8).

*Titers after accounting for estimated individual effects*

The data used in this paper contain sera collected from different individuals infected with the same variants. Even when these titers are grouped according to the infecting variant, one can see quite large variations in titer magnitudes whereas the qualitative appearance of the titer curve is similar (Fig. 1). Some of this is due to individual variation in the overall reactivity of sera collected from different individuals, for example related to the magnitude of the serological response. We model this behavior in the following way: Assume that $t_{ij}$ is the $\log_2$ titer measurement of antigen $i$ against serum $j$ from the serum group $J$. Then

$$t_{ij} = s_{iJ} + r_j + \varepsilon_{ij} (\mathrm{i} = 1, \dots \mathrm{n} \text{ and } \mathrm{j} = 1, \dots \mathrm{m})$$

where $s_{iJ}$ is the average $\log_2$ titer of antigen $i$ against the serum group $J$, $r_j$ is the serum reactivity bias of the serum $j$ and $\varepsilon_{ij}$ is the independently and normally distributed log titer noise. $n$ is the number of antigens and $m$ is the number of sera. For each serum group, parameters $s_{iJ}$, and $r_j$ are then found to maximize the sum of the likelihood of each measured log titer given the observed log titer $t_{ij}$. For the fitting procedure we used the function "estimate_sr_effects" in the "titertools" package, with parameter "sigma = 0.62", according to our estimates of the magnitude of measurement noise in a single measurement (fig. S8).

Note that antigenic cartography has its own method of dealing with serum reactivity bias. The formula for the target distance (section 'Antigenic cartography', below) indicates that for any serum, the antigen with the highest titer (usually the homologous antigen, but not always) is automatically adjusted to have 0 distance to that serum. This is true regardless of the value of the maximum titer and this also takes care of the individual serum reactivity in a way that is compatible with antigenic map making.

*Calculating fold-drop differences in vaccine sera*

When exploring differences between post-infection and different post-vaccination responses homologous to the same antigen, one might expect that the fold-drop in serum responses (compared to the homologous antigen) are similar in some way. A simple model one can use is to introduce an overall factor that relates the responses of sera from the two groups. In terms of the model in the previous section this means

$$s_{hJ} - s_{iJ} = d_J(s_{hK} - s_{iK}) = d_J \Delta {s_{hi}}^K$$

where $K$ and $J$ are two serum groups, $h$ is the homologous antigen to them, $i$ is any other antigen and $s_{xY}$ are as described in the previous section. Note that terms such as $s_{hJ} - s_{iJ}$ give the (noiseless) fold drop of titers of antigen $i$ from the titers of the antigen $h$ (homologous to $J$), measured against the serum group $J$. Then for a specific serum $j$ from the serum group $J$ we have

$$t_{hj} - t_{ij} = s_{hJ} - s_{iJ} + \varepsilon_{hj} - \varepsilon_{ij} = d_J \Delta s_{iK} + \varepsilon_{ijh}$$

where the last term is again normally distributed however with $\sqrt{2}$ times the standard deviation of the titer noise. An example would be $K$ = D614G post-infection serum group, $J$ = mRNA-1273 serum group, $h$ = D614G antigen and $i$ any other antigen (Figure 2). The term $d_J$ is the "factor" specific to the serum group $J$ which relates its responses to the response of the serum group $K$[1]. Note that taking difference of titers cancels out the serum response bias specific to the serum $j$.

When applying this model we compared the serum groups D614G, 4 weeks post 2x mRNA-1273, >3 months post 2x mRNA-1273 and 4 weeks post 3x mRNA-1273 (with shared homologous variant D614G) and compared the serum groups B.1.351 and 2x mRNA-1273.351 (with shared homologous variant B.1.351). In the D614G and

[1] This term obviously also depends on $K$ however we imagine that $K$ is a constant post-infection serum group and $J$ varies between different post-vaccination serum groups so by abuse of notation we only index this term with $J$.

B.1.351 post-infection groups we fixed $d_J$ as 1, estimating fold-change differences in the post-vaccination groups relative to the post-infection sera. For the fitting procedure we used the function "compare_crossreactivity_breadth" in the titertools package, with confidence intervals calculated for the parameters based on the highest density interval of 5000 sampling runs after 1000 warmup runs.

*Imputing censored titers*

For figures S11, S12 and S20 we impute non-detectable titers in order to not bias the distributions that are visualized. In these cases, censored titers (e.g. <20) are imputed using the function "impute_gmt_titers" from the "titertools" package. In these cases, censored titers are replaced with numeric titers drawn from a censored normal distribution with mean and standard deviation parameters that are estimated from the titer data as described above.

***Antigenic cartography***

Antigenic cartography is a multidimensional scaling method (MDS) (*64*) for the reduction of dimensions in large titer datasets in order to not only visualize large amounts of titer data in a lower-dimensional space but to also draw conclusions about the antigenic evolution of pathogens by studying the positions occupied by these viruses on the antigenic map.

In order to construct an antigenic map, one starts with a titer table where columns are sera and rows are antigens. Denoting the values of such a table by $t_{\mathrm{ij}}$ (where $i$ denotes the antigen index and $j$ the sera index), titer values represent a degree of dissimilarity of the $i^{\mathrm{th}}$ antigen to the $j^{\mathrm{th}}$ serum. We turn this dissimilarity score into a similarity score by the following transformation:

$$D_{ij} = \max_{\mathrm{i}}\left(\log_2\left(t_{\mathrm{ij}}\right)\right) - \log_2\left(t_{\mathrm{ij}}\right)$$

The $D_{ij}$ values are called the target distances. Antigenic cartography strives to find a multidimensional (usually 2 or 3 dimensions) placement of antigens and sera such that the Euclidean distance between antigens and sera conform as well as possible to the target distances. This is achieved by an optimization of the antigen and serum coordinates that minimize the stress function

$$\sum_{i,j} \left(D_{ij} - d_{ij}\right)^2$$

where $d_{ij}$ represents the Euclidean distance between the $i^{\mathrm{th}}$ antigen and the $j^{\mathrm{th}}$ serum points in the antigenic map. The minimization is achieved using the L-BFGS algorithm (*65*) to find the best cost-minimizing coordinates starting from a set of randomly determined initial conditions. Note unlike most common MDS problems, the similarity matrix is not square and might contain masked or censored values and therefore requires a slightly custom approach. In particular, if $t_{\mathrm{ij}}$ is a non-detectable titer, then the stress is computed as

$$\varphi(D_{ij} - d_{ij} + s)(D_{ij} - d_{ij} + s)^2$$

where s is a step size factor which is 1 for discrete data (and their means) such as those obtained from Haemagglutinin Inhibition assays, or 0 when the output data is continuous and measured from titer curves, and

$$\varphi(x) = \frac{1}{1 + 10^{-x}}$$

is a damping function whose role is to give more freedom to non-detectable titers to be optimized for a value less than the limit of detection of the assay.

We used the Racmacs package to compute the antigenic maps presented in this manuscript (*66*). The map was constructed using 1000 optimisations, dilution step size of 0, with the minimum column basis parameter set to "none". In order to allow for accurate coordination of serum positions in 2 dimensions, sera with less than 3 detectable titers were excluded from the maps. An exception was made for the BA.2 serum sample, which was included despite only having 2 detectable titers since it was the only serum of its group, although we note that the serum position for this serum will therefore be less constrained in its antigenic position.

*Assessing model fit*

We performed different analyses to ascertain that the two-dimensional antigenic map is a good representation of the titer data, and to ensure that the map is robust to noise from measurement errors.

First, to get an understanding of the measurement error of individual titrations, we analyzed a set of repeat titrations of 60 sera against 8 of the variants used in the assay, and found that the mean difference in $\log_2$ titers between repeated titrations was 0.209, with a standard deviation of 0.84, with measurement error being approximately normally distributed (fig. S8). Accounting for the systematic bias of titrations between repeats, and assuming a mean of 0 without it, resulted in a total standard deviation of 0.87 on the $\log_2$ scale. Since measurement error would be present for both the first and second repeat titration this would be consistent with noise per measurement with a standard deviation of $\sqrt{0.87^2/2} = 0.62$.

Subsequently, we investigated how well antigenic maps of different dimensions represent the measured patterns of serological reactivity. In our previous experience with other antigenically variable pathogens, the dimensionality of relationships can be lower (for example two dimensions in seasonal influenza and dengue virus (*12*, *67*)) or higher (for example three dimensions for A/H5N1 and swine influenza(*16*)), or may simply not be well represented in Euclidean space. Under cross-validation, a map of two dimensions represented the data best (fig. S9A). The overall arrangement of the variants and sera in three dimensions is similar to two dimensions (fig. S9B-C), with the exception of the BA.4/BA.5 variant, which in three dimensions occupies a position more distant from BA.1 and closer to B.1.617.2 and B.1.351 than visualized in two dimensions. The dimensional analysis was performed with Racmacs using the function “dimensionTestMap”.

*Assessing goodness of fit using fitted and measured titers*

Next, we assessed how well the two-dimensional map fits the measured titers. We first investigated whether the measured $\log_2$ titers correlated with the titers fitted in the antigenic map. Overall, the fitted and measured titers corresponded well (fig. S10). When comparing the mean differences of the fitted versus measured titers (mean: 0.04, standard deviation: 1.05, shown as a histogram in fig. S11), they are comparable to the estimated size of measurement error (fig. S8). This suggests that most of the residual error observed in the antigenic map fit of the data is consistent with measurement error, but that not all variation in the titers is captured.

We investigated whether any variant and serum group combinations exhibited consistently larger differences between fitted and measured titers (fig. S12). We find some variants showing differences in fitted versus measured titers in some serum groups, such as the B.1.617.2 variant having larger measured than fitted titers against the B.1.617.2 sera. Such differences do not necessarily result from antigenic dissimilarity but may also be related to changes in avidity of antibody binding (*32*). However, some of these systematic residual errors in the map fit will also be reflective of the fact that not all patterns of cross-reactivity between the serum groups can be captured in the constraints of the two-dimensional Euclidean space of the map. Overall though, we judge that the two-dimensional map is a reasonable representation of the broader scale underlying antigenic relationships among variants, providing a useful complement to raw titer analysis, and in some cases may compensate for systematic titer biases like that of P.1 that are otherwise spurious.

*Assessing the robustness of variant and serum positions to measurement error and presence/absence of variants and serum groups*

We performed three experiments to determine the uncertainty of the positions of variants and sera in the antigenic map. First, we estimated how titration measurement error and variability in antigen reactivity would affect conclusions about antigenic map positions (fig. S13). We performed a “smooth” bootstrap with normally distributed noise added to the titer measurements and / or to antigen reactivity. Noise added to the titer measurement had a standard deviation of 0.62, in keeping with the standard deviation of repeat titrations shown in fig. S8, and random noise for antigen reactivity had a standard deviation of 0.4, indicating that systematic biases in titers like that suspected for P.1 may also be present to some degree for other strains in the assay. We performed 500 repeats of adding random noise (using the Racmacs function bootstrapMap, method = “noisy”), and summarized the variant positions by showing the area in which 68% (one standard deviation) of the positional variation of a serum or antigen is captured. In general, we found the antigen and serum positions are robust to both the random and systematic noise added, without affecting the main conclusions about the antigenic relationships between the variants included.

Next, we assessed whether the positions of the variants are robust to the exclusion of each variant in turn. We find that positions of variants are robust to the exclusion of variants (fig. S14), with the small movement observed with the exclusion of D614G and B.1.351. We then investigated whether positions of variants are robust to the exclusion of each serum group in turn (fig. S15). Overall, the map was robust to the exclusion of single serum groups, with the greatest movement observed when excluding the 2x mRNA-1273.351 vaccine serum group and the B.1.617.2 and BA.1 convalescent serum groups. We also included the outlier sera identified in fig. S1, and found little difference in map topology (fig. S17).

We also evaluated the robustness of map positions on particular titrations by performing 500 “Bayesian bootstrap” repeats, where in each repeat, titers were reweighted according to a dirichlet distribution . This is statistically similar to bootstrapping by taking random samples with replacement but ensures that titers are never fully removed from the dataset, which can happen when sampling with replacement and lead to under constrained antigenic map solutions in some of the bootstrap repeats. We performed this reweighting on both variants and sera combined, as well as antigens and sera separately, using the Racmacs function bootstrapMap, method = “bayesian” (fig. S18). Map positions were largely robust to this reweighted sampling of titers.

We also show that the error is approximately equally distributed between all variants and sera (fig. S19A). Positions of variants and sera are also robust when allowing the variant and sera to move in the map while not increasing the overall map RMSE between map and table distance by more than one unit (fig. S19B).

*Assessing predictive power of the antigenic map (cross-validation)*

To investigate whether the antigenic map is able to predict missing titers, we performed 500 cross-validation replicates, where in each replicate we removed 10% of all titrations, and predicted the missing titers from the resulting map. On the $\log_2$ scale, where difference of 1 represents a predicted titer that is 2-fold higher than the measured titer and -1 represents a titer that is 2-fold lower, the mean difference between predicted and measured titer was 0.53 with a standard deviation of 1.61 (fig. S20). We also considered the difference between predicted and measured titers split by variant and serum group (fig. S21). As for fig. S11 and fig. S12, these results imply that not all patterns of cross-reactivity between the serum groups can be captured in the constraints of the two-dimensional Euclidean space of the map, but that overall the two-dimensional relationships are largely predictive of titers among sera and variants.

***Construction of the Antibody Landscapes***

The construction of the antibody landscapes differs from the methodology used in previous publications (*32*, *68*), since each serum represents a likely first infection and was used to construct the map itself with explicit assumptions about how reactivity should be distributed. Given a serum with coordinates $(x_m, y_m)$ let $t_m$ be its maximum $\log_2$ titer/10. Then $(x_m, y_m, t_m)$ represents the highest point of the antibody landscape for this serum. Given any other point $p = (x, y)$ on the two-dimensional base space, the third coordinate $t$ of the surface at this point is given by

$$t = t_m - d(p, p_m)$$

where $d(p, p_m)$ is the Euclidean distance between $p$ and $p_m$. In particular, the serum point and maximum titer for a given serum describe how reactivity would be expected to vary in a cone-like fashion across antigenic space, with its apex at the serum point at a height equal to the maximum measured $\log_2$ titer/10 and diminishing at a rate of a two-fold reduction in titer with each antigenic unit on the map. The serum group averages shown in Fig. 4 therefore simply represent an average of all these individual slope = 1 cone-like responses for the relevant sera, displaying the GMT of all the individual titer predictions for the group.

In this case, since samples were collected post a presumed single exposure, individual landscapes simply visualize the slope 1 cone-like model of reactivity that is fit when constructing the antigenic map, rather than refitting the same data with a nonparametric modeling approach that can also be applied.

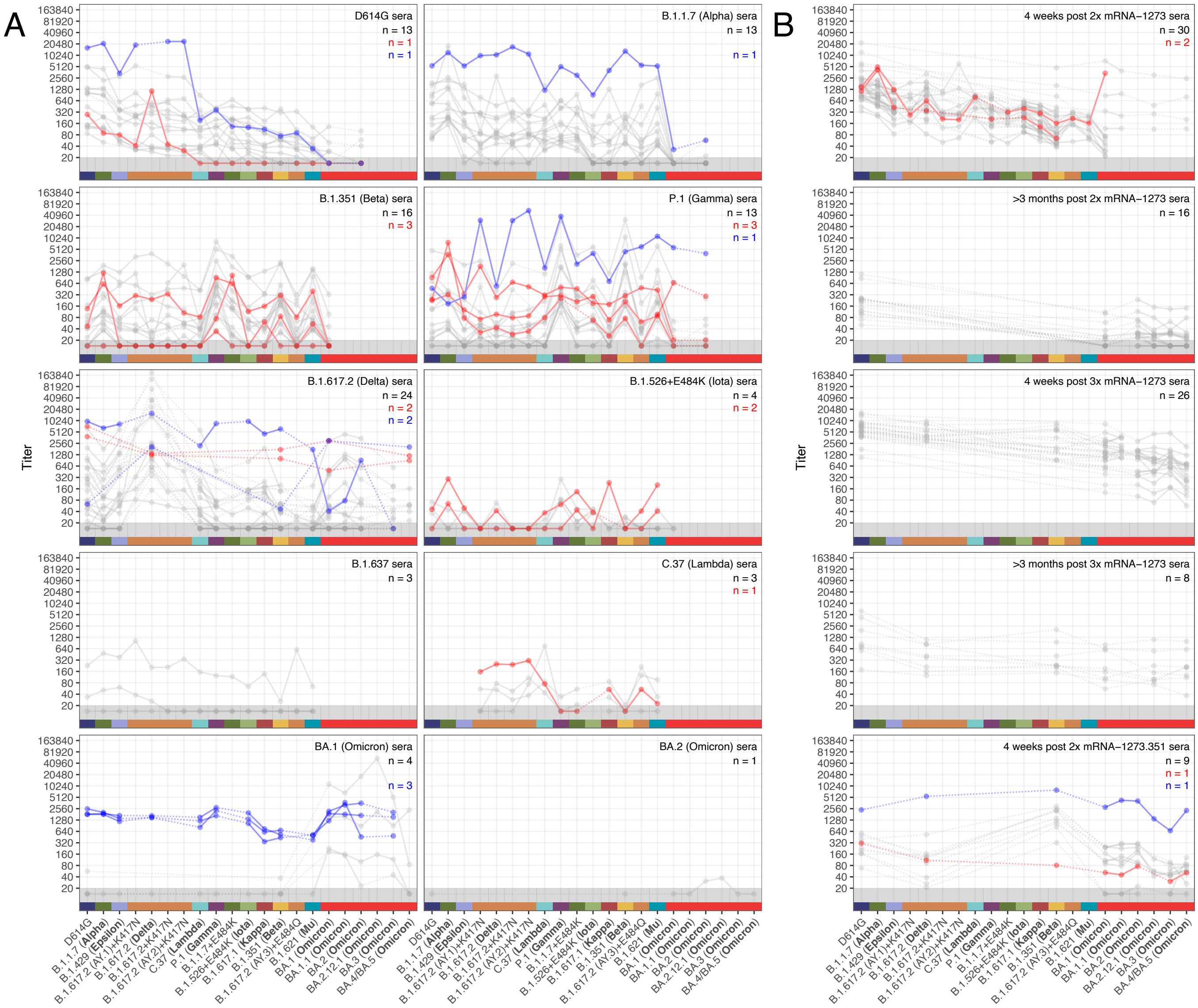


**Figure S1: Complete set of sera highlighted as possible non-primary exposures.** As with Fig.1, variants are ordered according to GMT in D614G sera (top-left), which allows for direct comparison in a column. Fainter individual lines and solid points show individual titer traces. Lines relating to subjects that met exclusion criteria based on presence of a titer that was >2-fold higher than the titer against the homologous variant for their serum group are shown highlighted in red. Lines relating to subjects that were excluded based on visual inspection of titer patterns are shown highlighted in blue. Numbers relating to the total number of non-excluded sera and sera excluded by each approach for each serum group are shown in the top right corner of each plot. Points in the gray region at the bottom of the plots show titers and GMTs that fell below the detection threshold of 20. GMT for data including non-detectable titers was calculated as described in Materials and Methods, ‘Titer Analyses’ section.

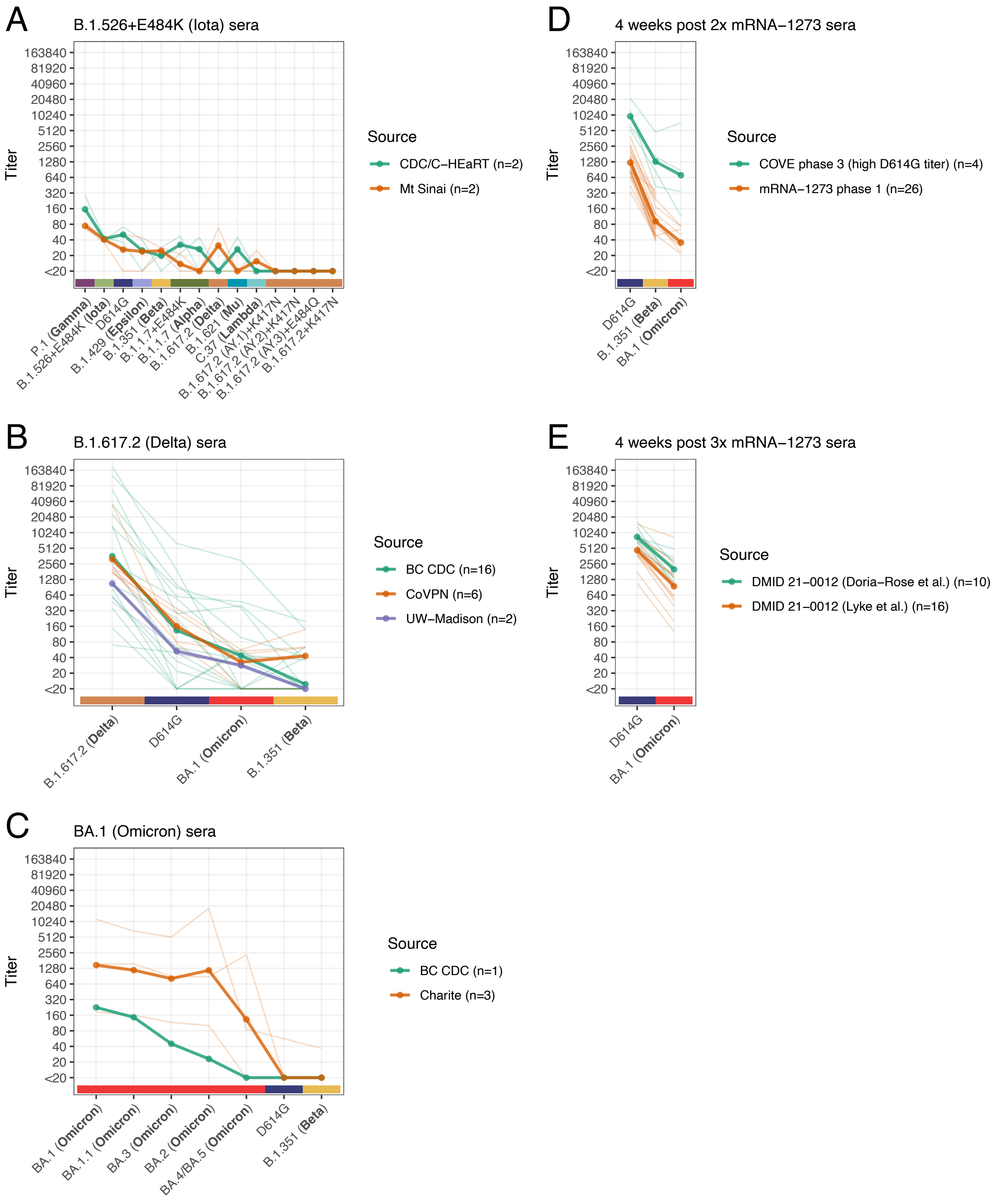


**Figure S2: Comparison of titers between samples taken from different cohorts or agencies.** Each panel shows data from the five serum groups where samples came from a mixture of cohorts or agencies (after removal of outlier sera shown in fig. S1). Fainter lines show the pattern of titers for each individual sample, colored corresponding to the source of each sample. Bolder colored lines and points show the GMT from all the samples from each source, calculated as described in Materials and Methods. In each panel, the variants included are those that were jointly titrated against samples from each of the sources and variants are ordered by decreasing mean GMT.

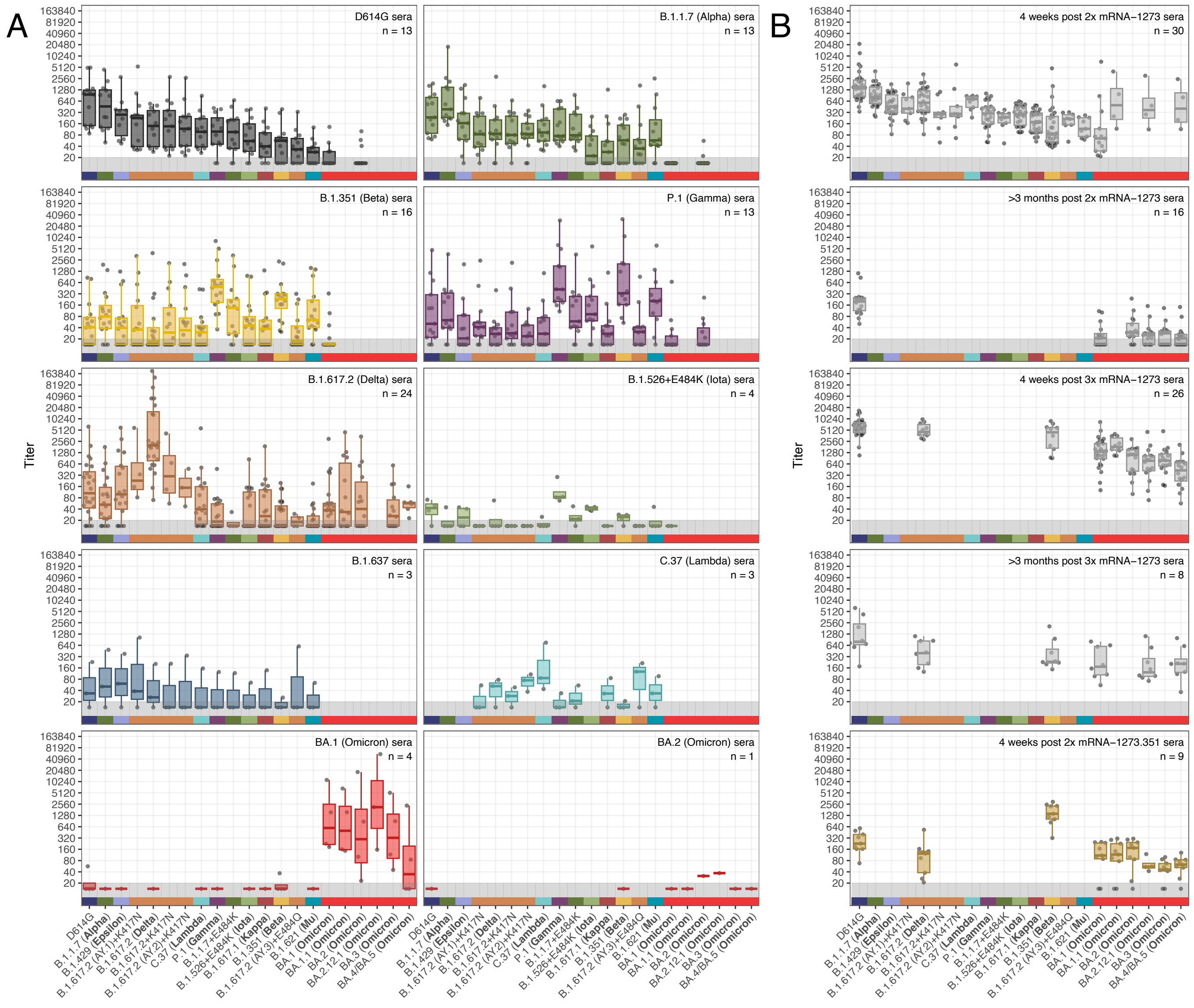


**Figure S3: Neutralization of lentivirus pseudotypes encoding different SARS-CoV-2 spike proteins by sera from patients vaccinated or infected with different SARS-CoV-2 variants.** The box plot indicates the median and 25th and 75th percentile. Black circles show the individual titers. Variants are ordered according to GMT in D614G sera (top-left), which allows for direct comparison in a column. The gray region designates titers that fell below the limit of detection of 20. Titers shown correspond to the line plot in the main manuscript Fig. 1.

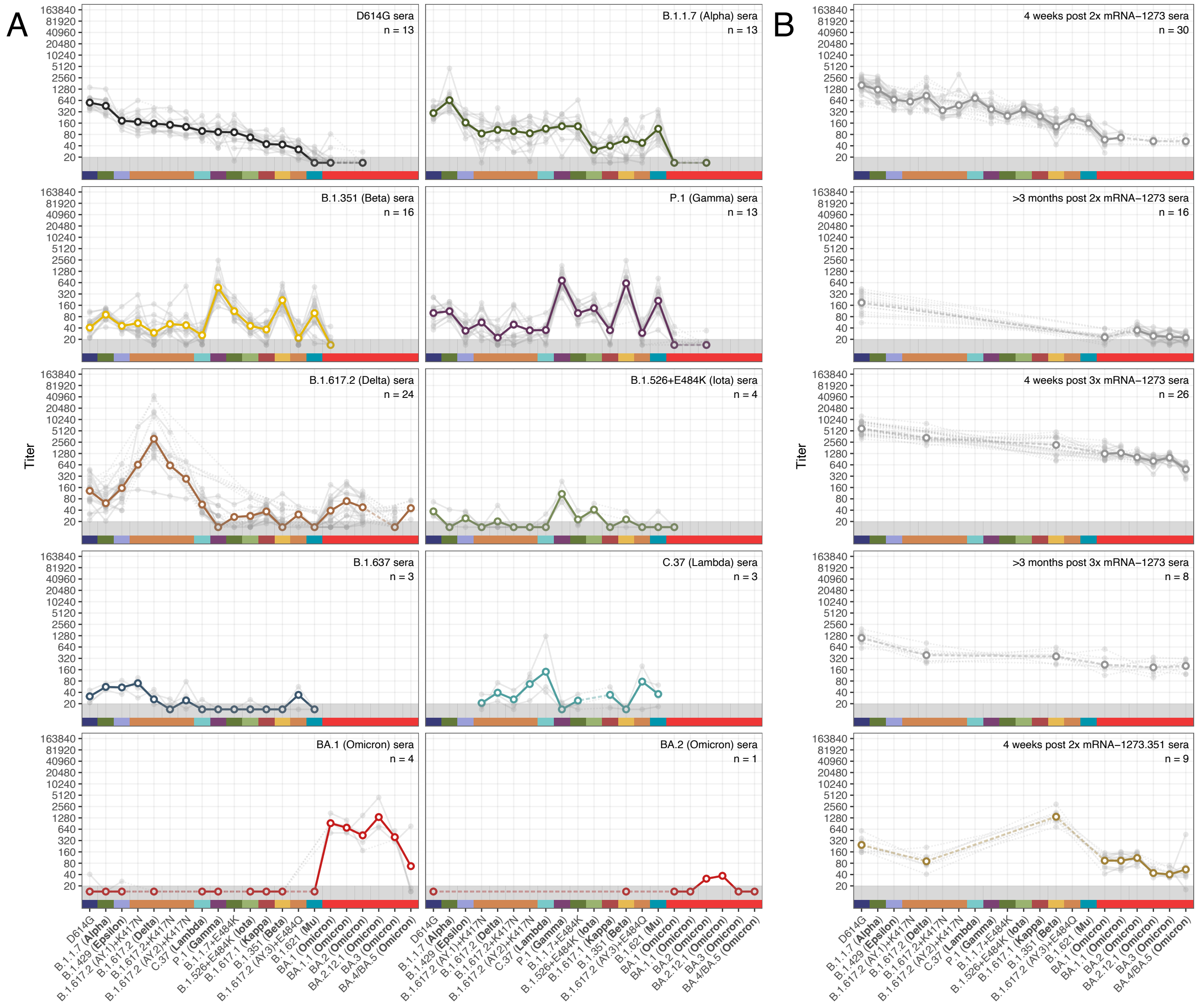


**Figure S4: Neutralization of lentivirus pseudotypes encoding different SARS-CoV-2 spike proteins by sera from patients vaccinated or infected with different SARS-CoV-2 variants after accounting for individual effects.** Individual effects were accounted for as described in Materials and Methods, ‘Titers after accounting for estimated individual effects’. Variants are ordered according to GMT in D614G sera (top-left), which allows for direct comparison in a column. White outlined points and the dense line shows the GMT, fainter individual lines and solid points show individual titer traces. Points in the gray region at the bottom of the plots show titers and GMTs that fell below the detection threshold of 20.

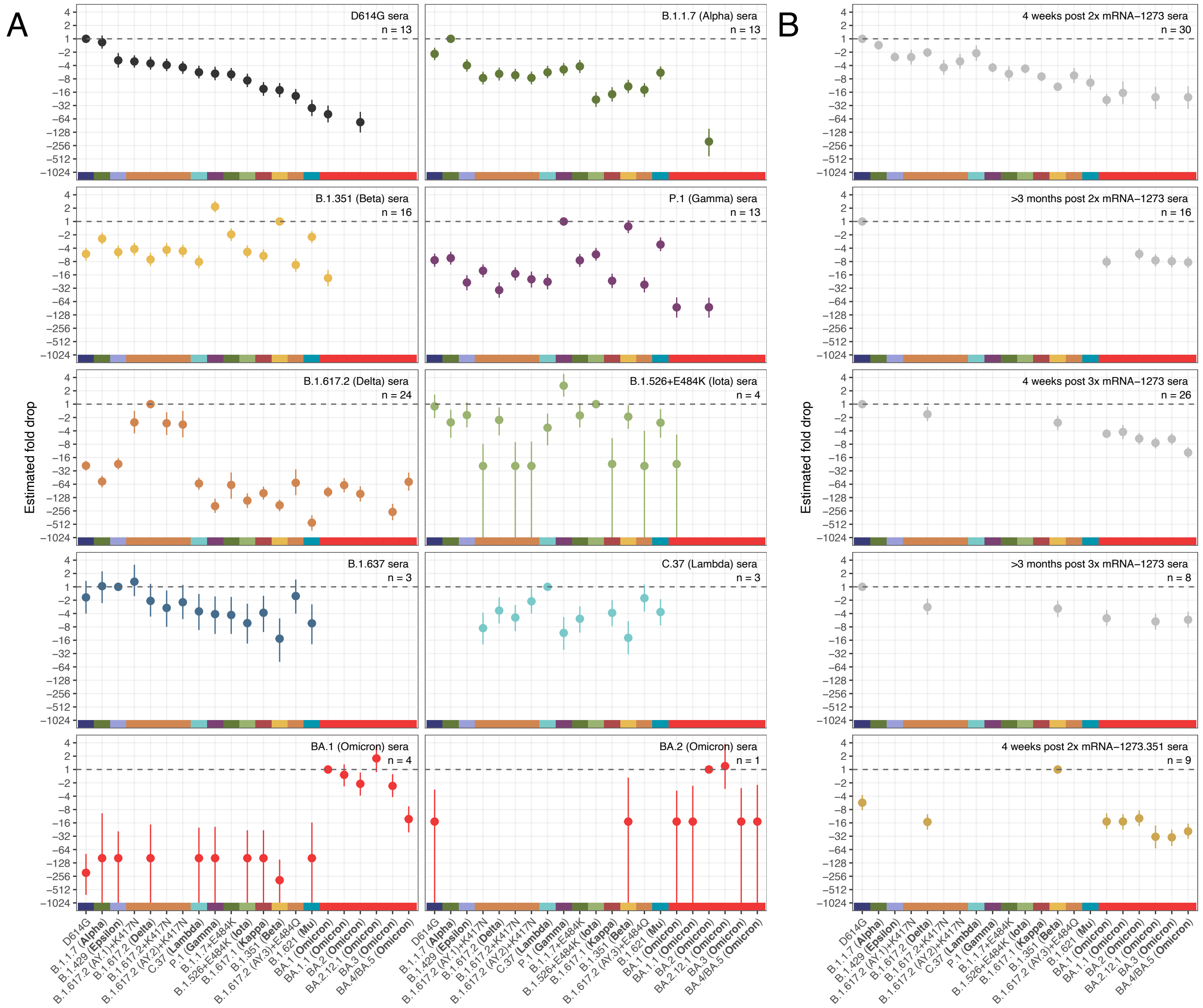


**Figure S5: Estimates of fold-change compared to the homologous variant.** Points show the point estimate for the fold-change difference while lines show the 95% HDI for the estimate. Fold drops and confidence intervals were calculated as described in Materials and Methods, "Titer analyses". As for figure 1, variants are order according to GMT estimates for the D614G sera.

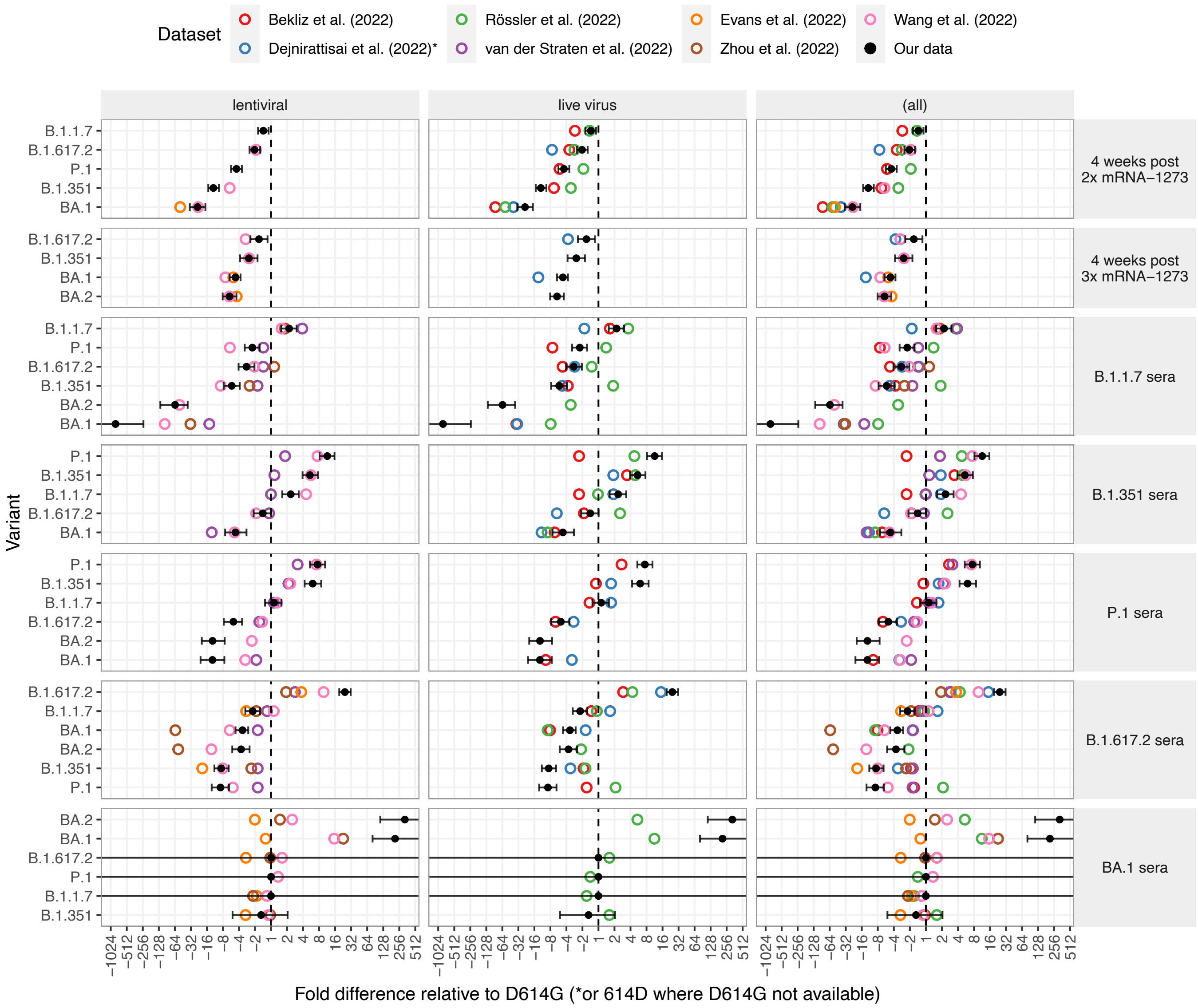


**Figure S6: Comparison of serum group fold-differences with results reported from other studies.** The $\log_2$ fold changes from D614G to other variants were calculated from the GMTs given in other studies (open colored circles) and the current study (solid black circle), according to the serum group indicated in each row. Fold differences shown are for the variant on the y axis relative to the D614G titer or, in the case of Dejnirattisai et al., relative to 614D. Black error bars indicate the 95% HDI based on our own data, calculated as described in Materials and Methods, “Titer Analyses”. Comparative study results are split by “lentiviral” and “live virus” type in the first and second columns and combined in the third. In more detail, assay types were: Bekliz et al.; Live-virus in Vero-E6, Dejnirattisai et al.; Live-virus in Vero (614D not D614G), Evans et al. (2022); Lentiviral pseudotype in 293T-ACE2, Rössler et al. (2022); Live-virus in Vero ACE2/TMPRSS2, van der Straten et al. (2022); Lentiviral pseudotype in 293T-ACE2, Wang et al. (2022); Lentiviral pseudotype in 293T-ACE2/TMPRSS2, Zhou et al. (2022); Lentiviral pseudotype in 293T-ACE2.

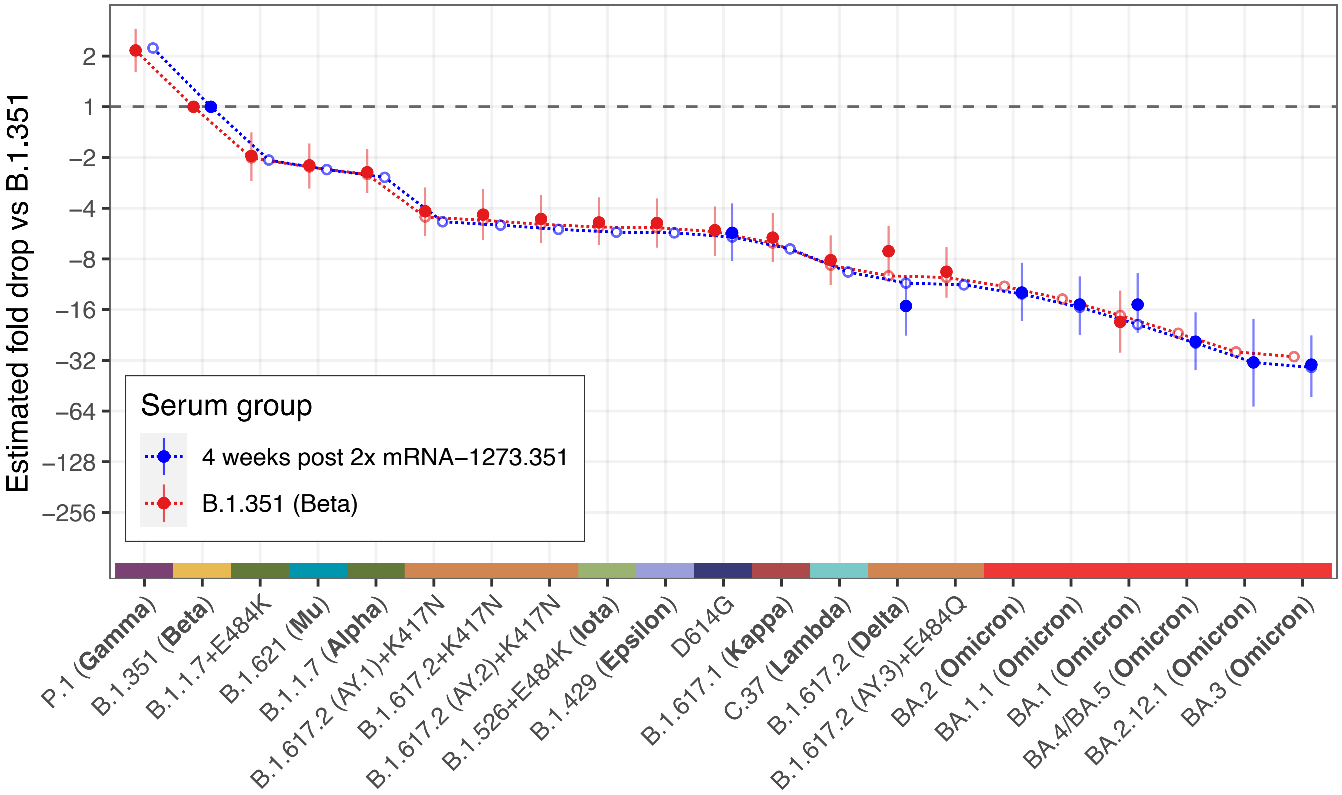


**Figure S7: Comparison of the mean fold drop relative to the homologous B.1.351 variant in B.1.351 convalescent sera and 4 weeks post 2x mRNA-1273.351 vaccination sera.** Solid points show the estimate for the mean fold drop compared to the titer for B.1.351, while lines represent the 95% HDI for this estimate. The points for B.1.351 to the left of the plot represents the homologous virus against which fold-change for other strains was compared and are therefore fixed at 1. Dotted lines and outline circles show estimates based on a model that assumes a shared overall pattern of fold-drops but estimates “slope” differences in the rate of reactivity drop-off seen in the 2 serum groups, as described in Materials and Methods, “Calculating fold-drop differences in vaccine sera”. To aid comparison, points and lines for each of the serum groups have some offset in the x-axis.

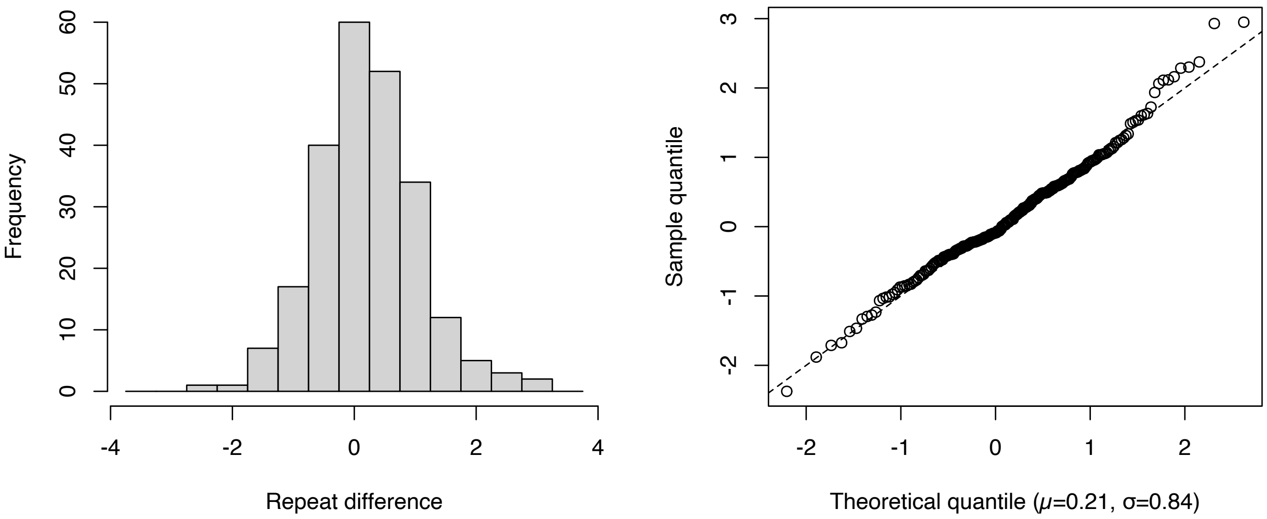


**Figure S8: RMSE from repeat titrations.** The left figure shows the histogram of differences between repeat titrations, while the quantile-quantile plot on the right indicates that measurement error is approximately normally distributed. The mean difference between repeated titrations was 0.21 [95% CI 0.1, 0.32], with a standard deviation of 0.84. Assuming instead a mean of 0 to account for the systematic bias of titrations between repeats, the standard deviation for measurement error is 0.87 on the $\log_2$ scale. Accounting for the presence of measurement error in both the first and second titration the standard deviation of noise per measurement is $\sqrt{0.87^2/2} = 0.62$.

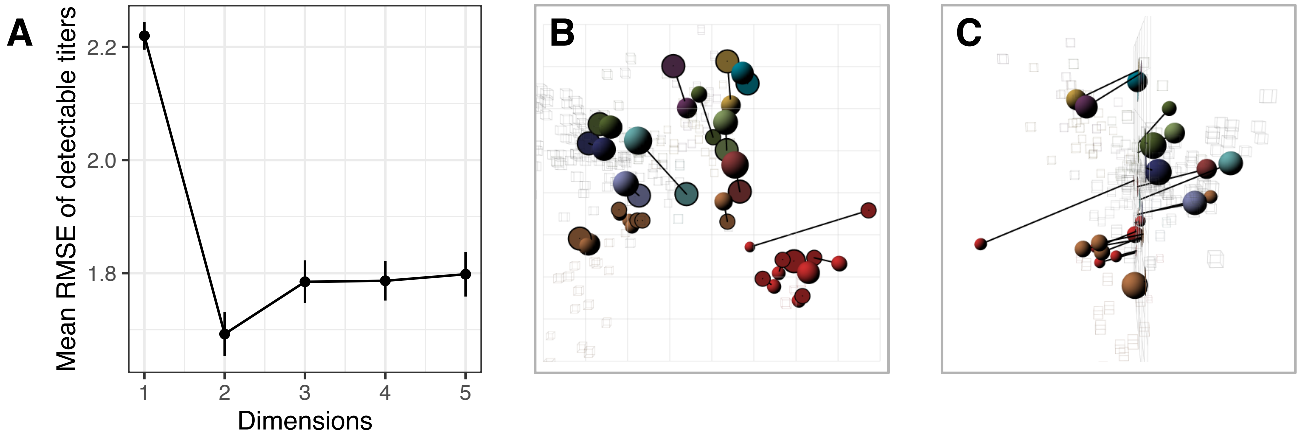


**Figure S9: Dimensionality tests and 3D map.** A) RMSE of detectable titers in 1 to 5 dimensions compared to known titers. Per dimension, 200 repeats were performed wherein a map was constructed from 90% of all titers. For each run, the RMSE is calculated by comparing the titers predicted from the map against the known titers on the $\log_2$ scale. No reduction in error is achieved when using more than two dimensions. B, C) The map optimized in 3 dimensions, with arrows pointing to where each variant is placed in two dimensions. B) 'Side' view. C) 'Front' view. The red antigen on the left is BA.4/BA.5 which is better fitted in 3 dimensions.

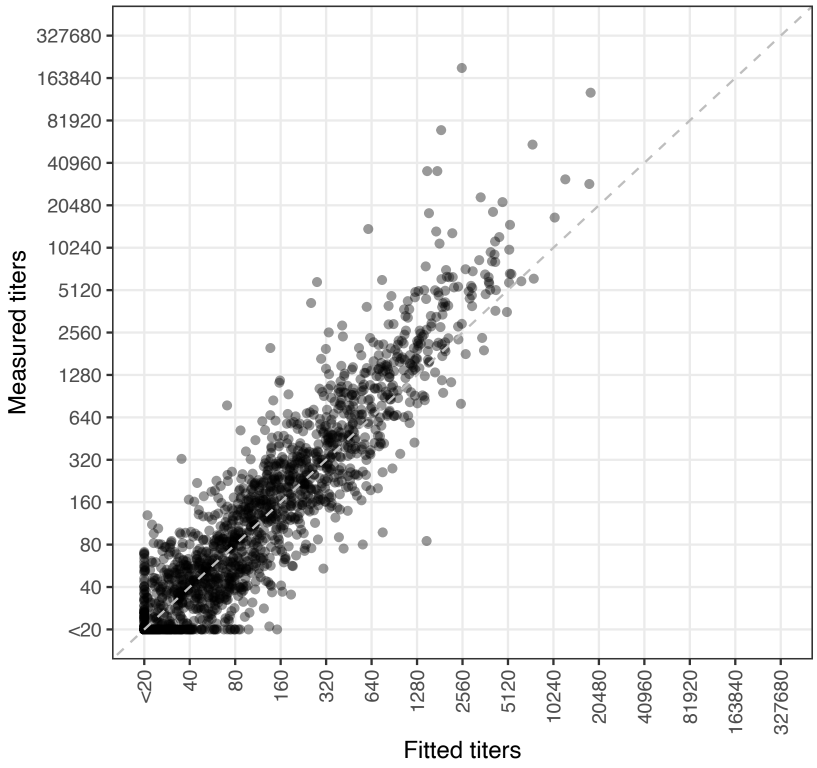


**Figure S10: Scatter plot of fitted titers against measured titers.** Titers are plotted on the $log_2$ scale. Fitted titers are determined from the distances in the antigenic map. The dashed gray line is the line of equality.

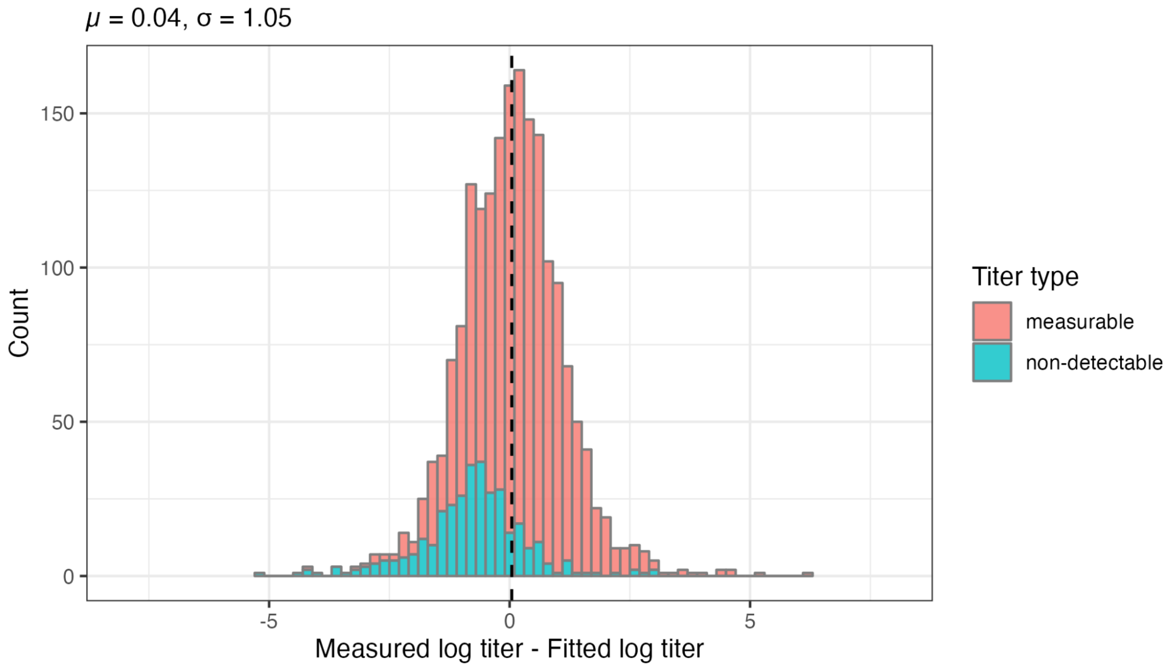


**Figure S11: Histogram of differences between fitted and measured $\log_2$ titers.** For non-detectable measured titers, differences between fitted and measured titers were imputed as described in Materials and Methods and are indicated in blue. Differences for detectable measured titers are indicated in red. The black dashed line indicates the mean.

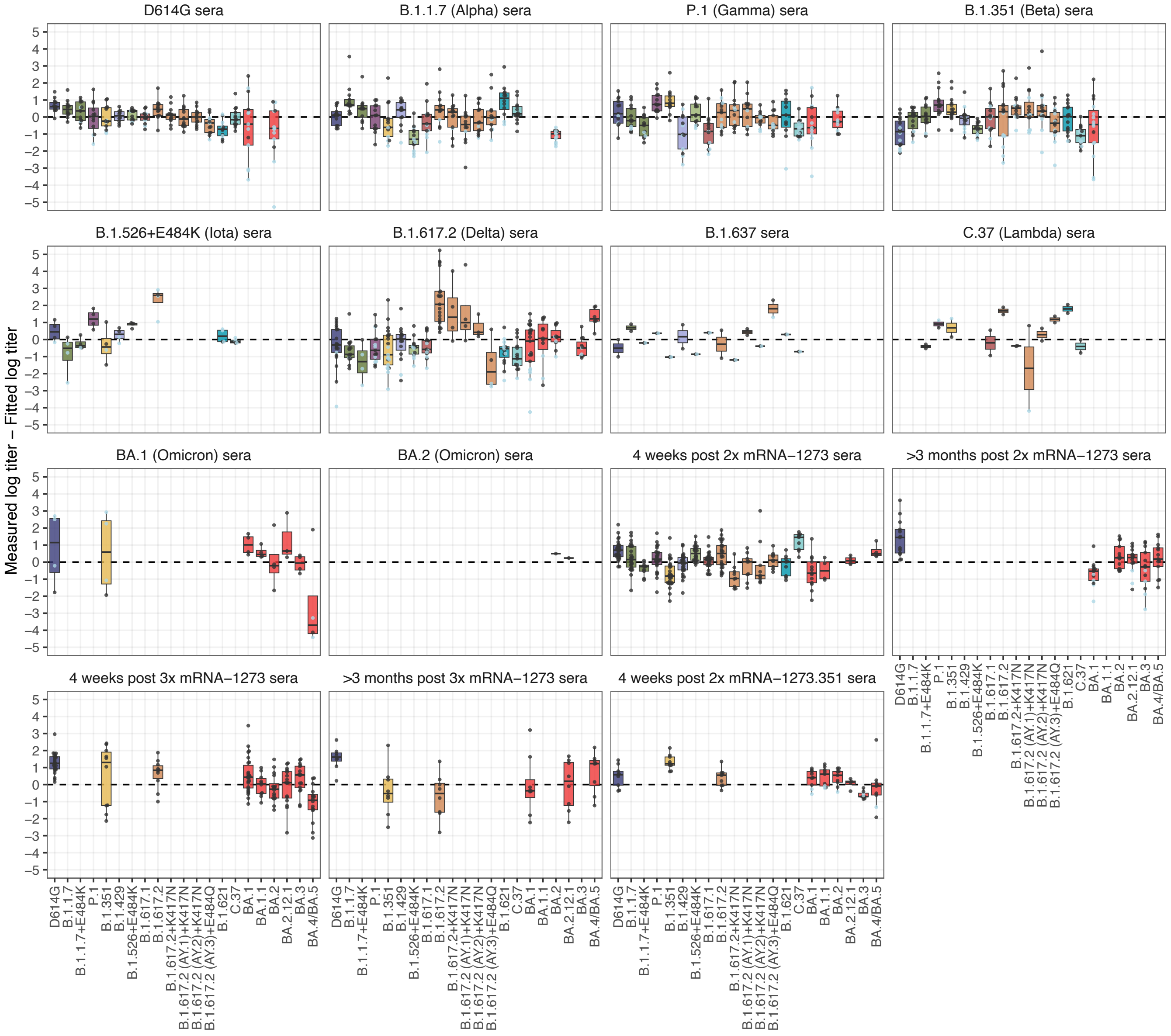


**Figure S12: Measured minus fitted $log_2$ titers split by serum group and variant.** The difference between fitted and measured $log_2$ titers is shown split by variant and serum group. Circles indicate the residual for each serum titration. Residuals for where the measured titers are non-detectable are shown in light blue and were imputed, residuals where the measured titers were detectable in black. The box plot indicates the median and 25$^{th}$ and 75$^{th}$ percentile.

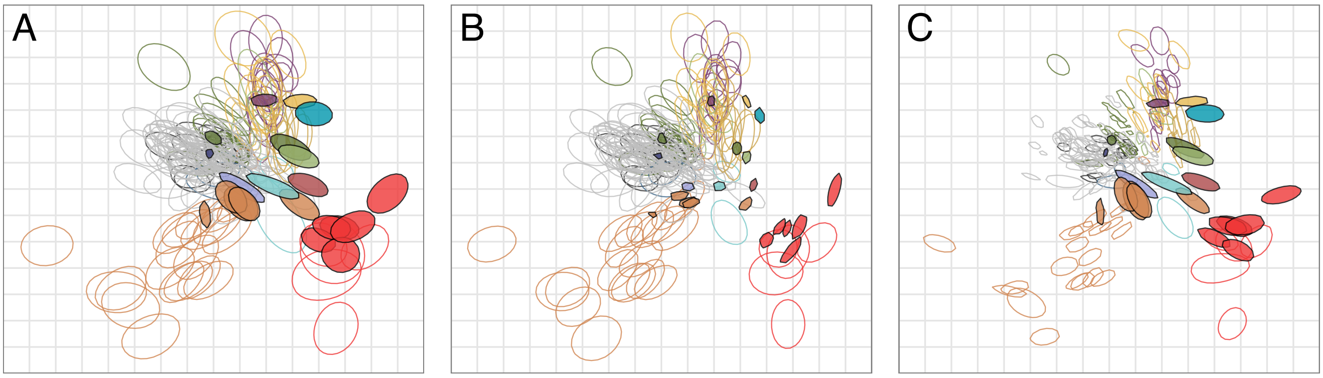


**Figure S13: Effect of the uncertainty in titers and variant reactivity for the antigenic map using "smooth" bootstrap.** 500 bootstrap repeats were performed with 500 optimizations per repeat. A) Bootstrap with noise added to titers and antigen reactivity. B) Bootstrap with noise added to titers. C) Bootstrap with noise added to antigen reactivity. The noise added to the titers was normally distributed and had a standard deviation of 0.62, the noise added to the antigen reactivity a standard deviation of 0.4. Each colored region indicates the area in which 68% (one standard deviation) of the positional variation of a serum or variant position is captured. Triangles indicate the positions of variants and sera that are outside the area of the plot. For color correspondence of variants and sera, refer to Fig. 3.

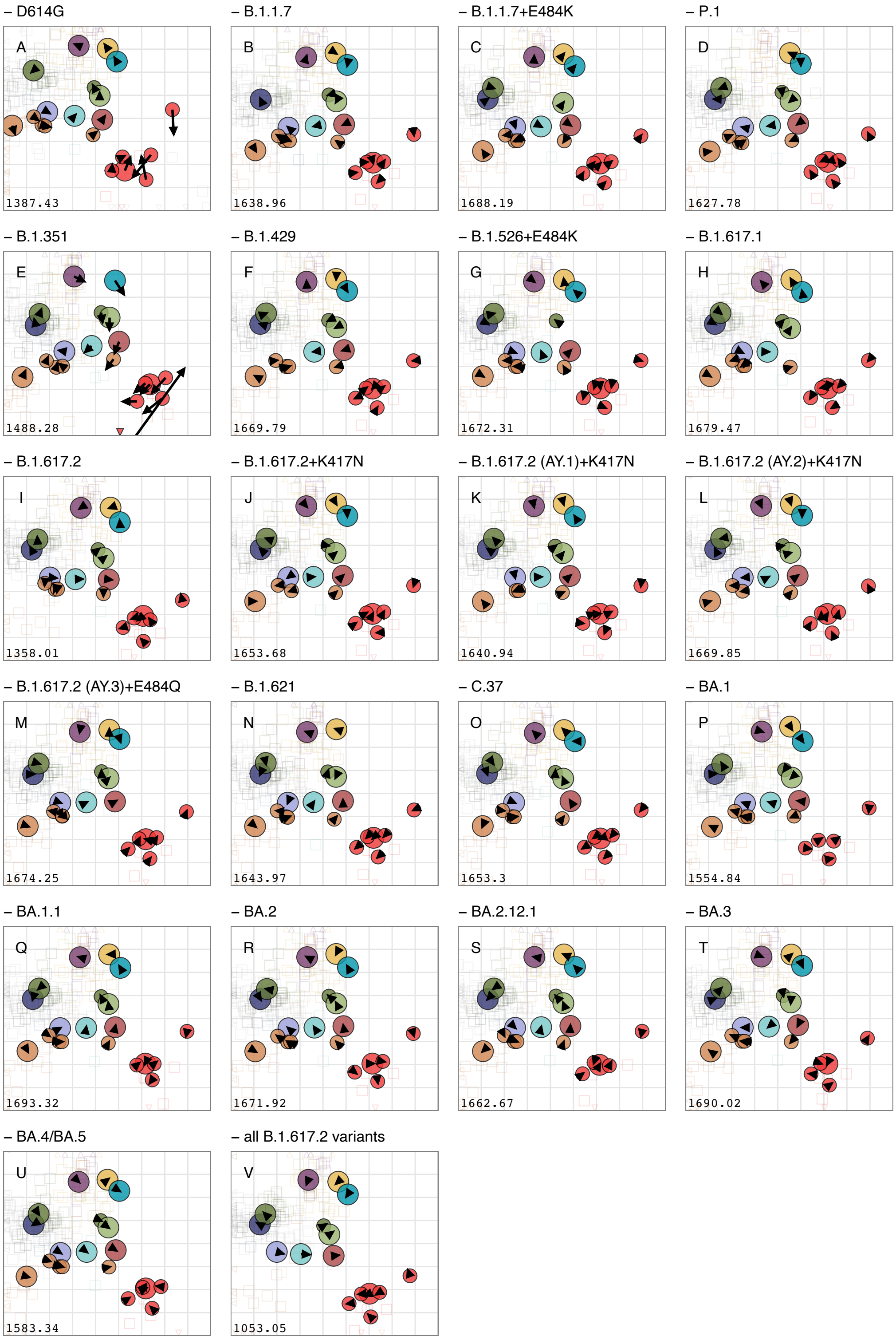


**Figure S14: Effect of removing variants on map topology.** Each variant was removed in turn and the map re-optimized. Arrows point to the positions of the variants in the map with all variants (shown in Fig. 3). Triangles indicate the positions of sera that are outside the area of the plot. Removed variants: A) D614G, B) B.1.1.7, C) B.1.1.7+E484K, D) P.1, E) B.1.351, F) B.1.429, G) B.1.526+E484K, H) B.1.617.1, I) B.1.617.2, J) B.1.617.2+K417N, K) B.1.617.2 (AY.1)+K417N, L) B.1.617.2 (AY.2)+K417N, M) B.1.617.2 (AY.3)+E484Q, N) B.1.621, O) C.37, P) BA.1, Q) BA.1.1, R) BA.2, S) BA.2.12.1, T) BA.3, U) BA.4/BA.5, V) All B.1.617.2 variants. For color correspondence of variants and sera, refer to Fig. 3.

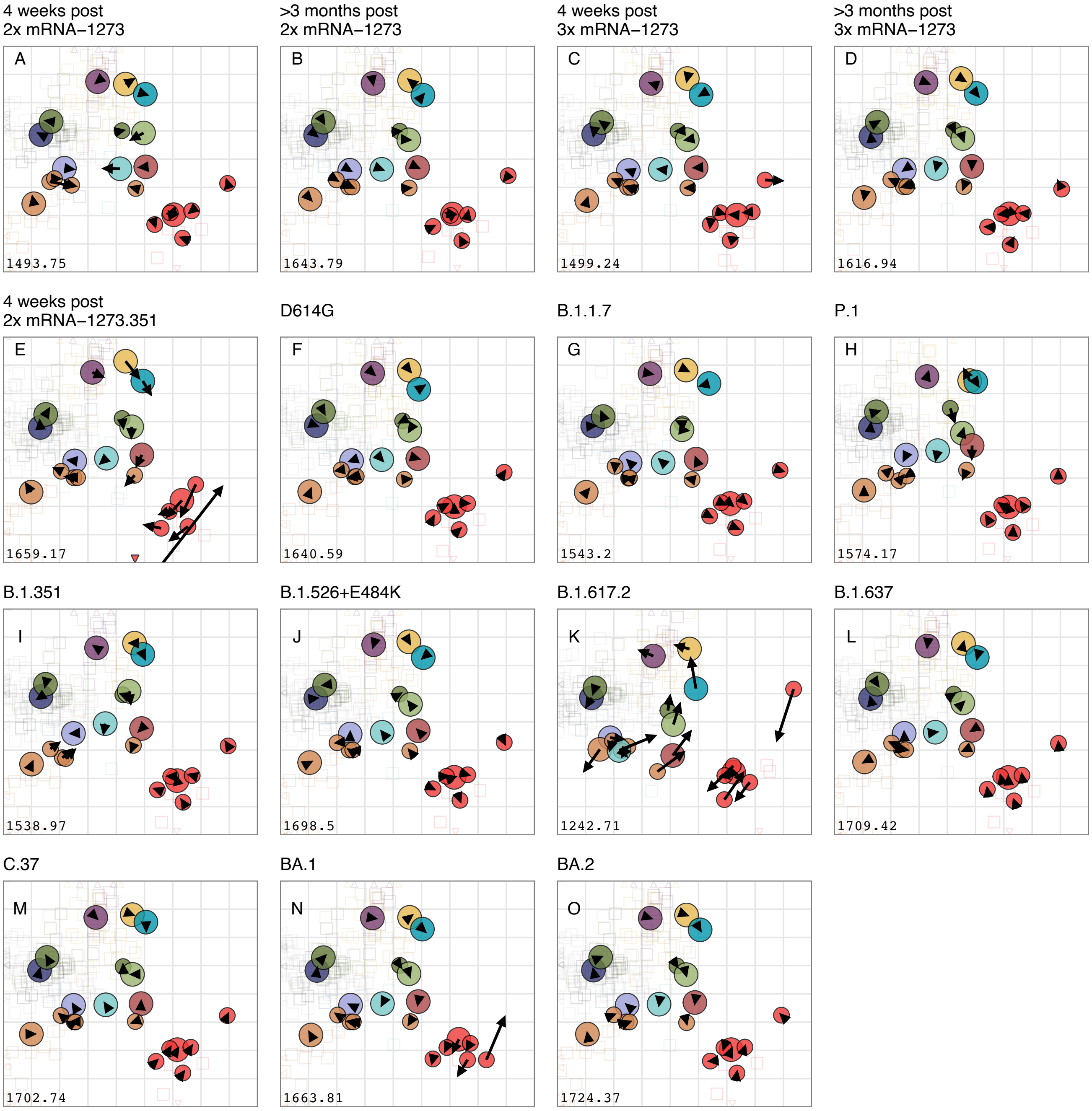


**Figure S15: Effect of removing different serum groups on map topology.** Each serum group was removed in turn and the map re-optimized; the removed serum group is given in the panel title. Arrows point to the positions of the variants in the map with all variants (shown in Fig. 3A), a short arrow thus indicates little change in position of a variant. Triangles indicate the positions of sera that are outside the area of the plot. For color correspondence of variants and sera, refer to Fig. 3.

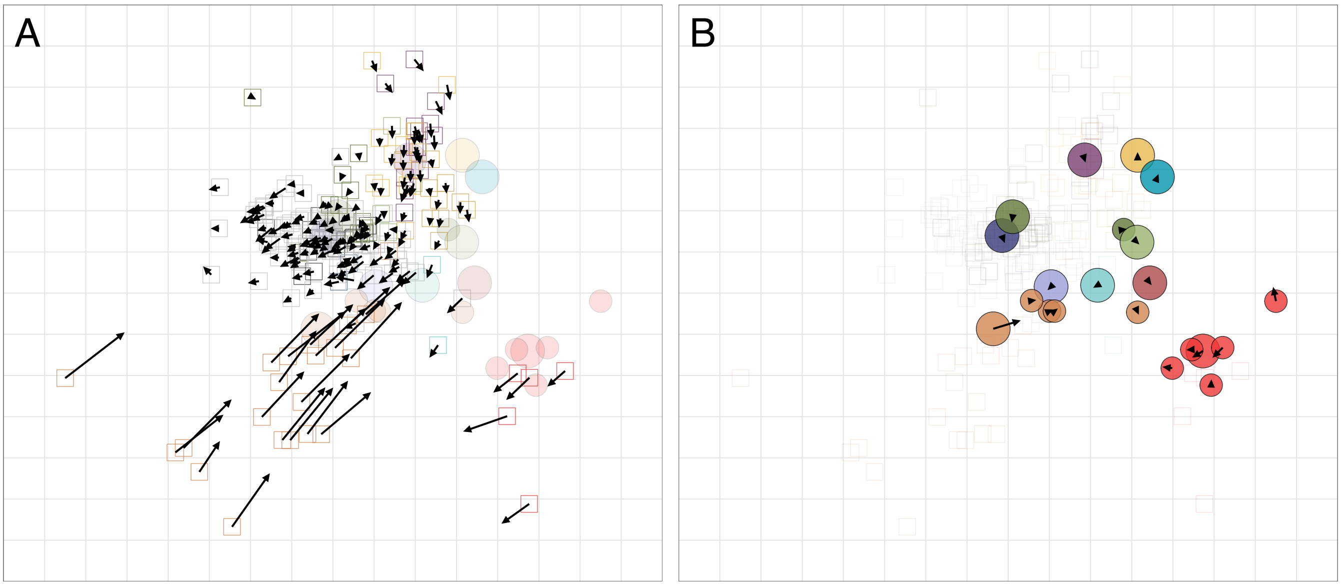


**Figure S16: Effect of reducing B.1.617.2 homologous B.1.617.2 titers by four-fold on map topology.** The antigenic map was re-optimized after the reduction in B.1.617.2 titers. In each panel, the map shown in Fig. 3A is reproduced with arrows showing how the position of sera (panel A) and antigens (panel B) varies compared to the titer-adjusted map. Predominant effects of the titer adjustment are to move B.1.617.2 serum positions inwards towards B.1.617.2 and to move the B.1.617.2 variant inwards towards the three B.1.617.2 + K417N variants and B.1.429 (Epsilon).

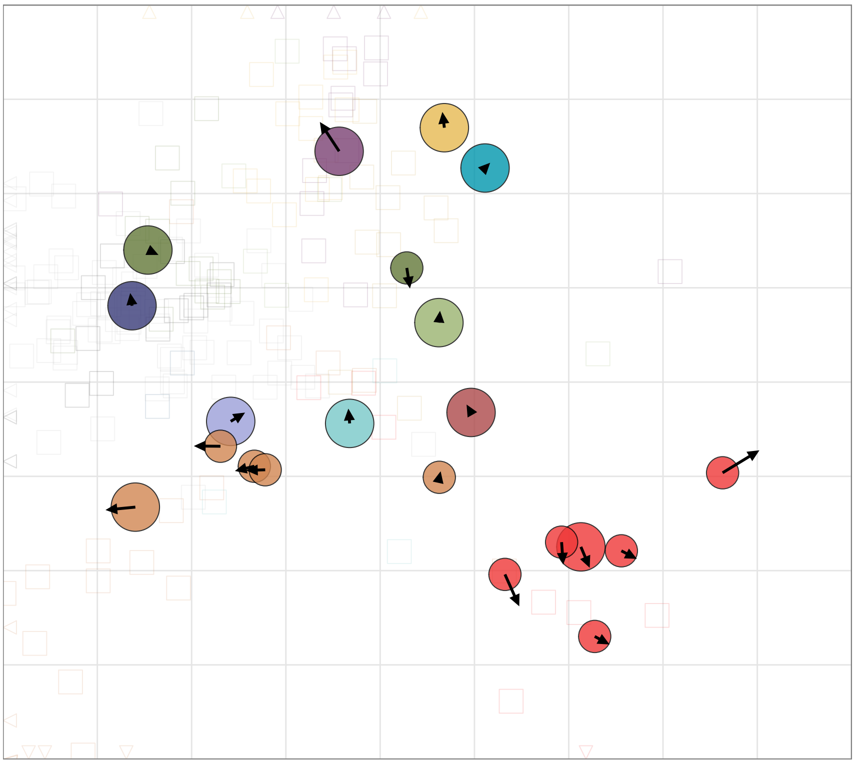

**Figure S17: Assessing the effect of outlier sera removal on map topology.** Map, including outlier sera that were identified as described in section ‘Excluding outlier sera’ and shown in fig. S1. Arrows point to the positions of the variants in the map shown in figure 3A.

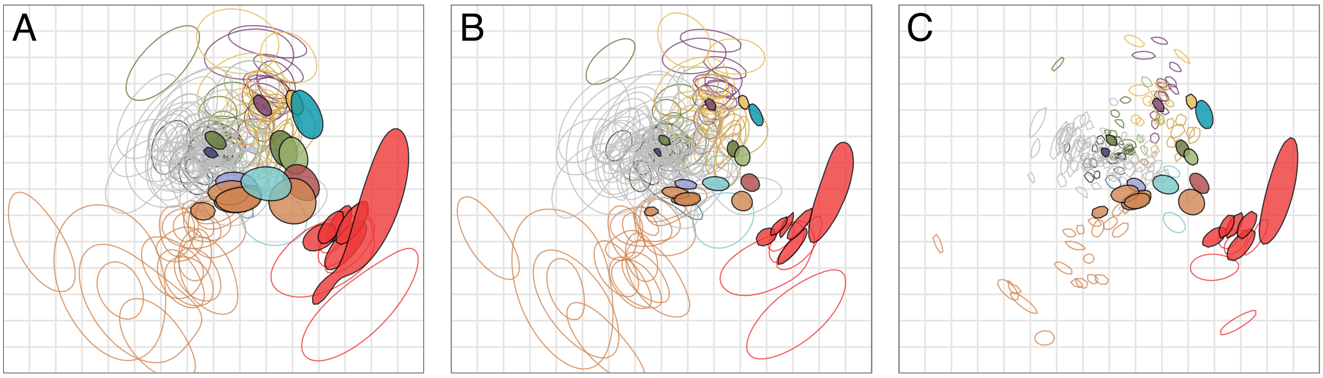


**Figure S18: Assessing the robustness of the antigenic map to different titer measurements.** 1000 bootstrap repeats were performed where a random weighted sample of titers was taken. A) Random weighting is applied by titer. B) Random weighting is applied by variant. C) Random weighting is applied by serum. Each colored region indicates the area in which 68% (one standard deviation) of the positional variation of a serum or variant position is captured. For color correspondence of variants and sera, refer to Fig. 3.

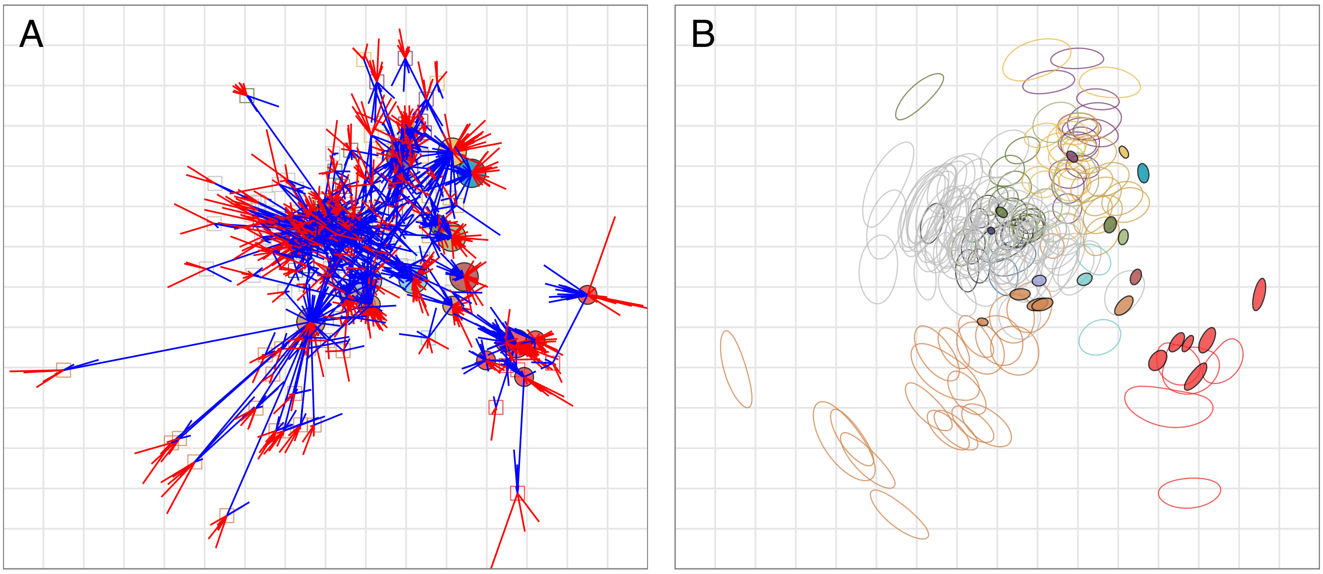


**Figure S19: Maps showing the uncertainty of the placements of variants and sera.** A) Antigenic map with error lines. For each titration, error lines are drawn between the variants and the serum that show the target distance between the variant and serum. Blue lines indicate a target distance shorter than the distance between the variant and serum in the map, red lines indicate that the distance on the map is shorter than the distance in the table. B) Triangulation blobs. The colored region for each serum and variant point indicates the area which the variant or serum can occupy without the stress of the map increasing by more than one unit. For color correspondence of variants and sera, refer to Fig. 3.

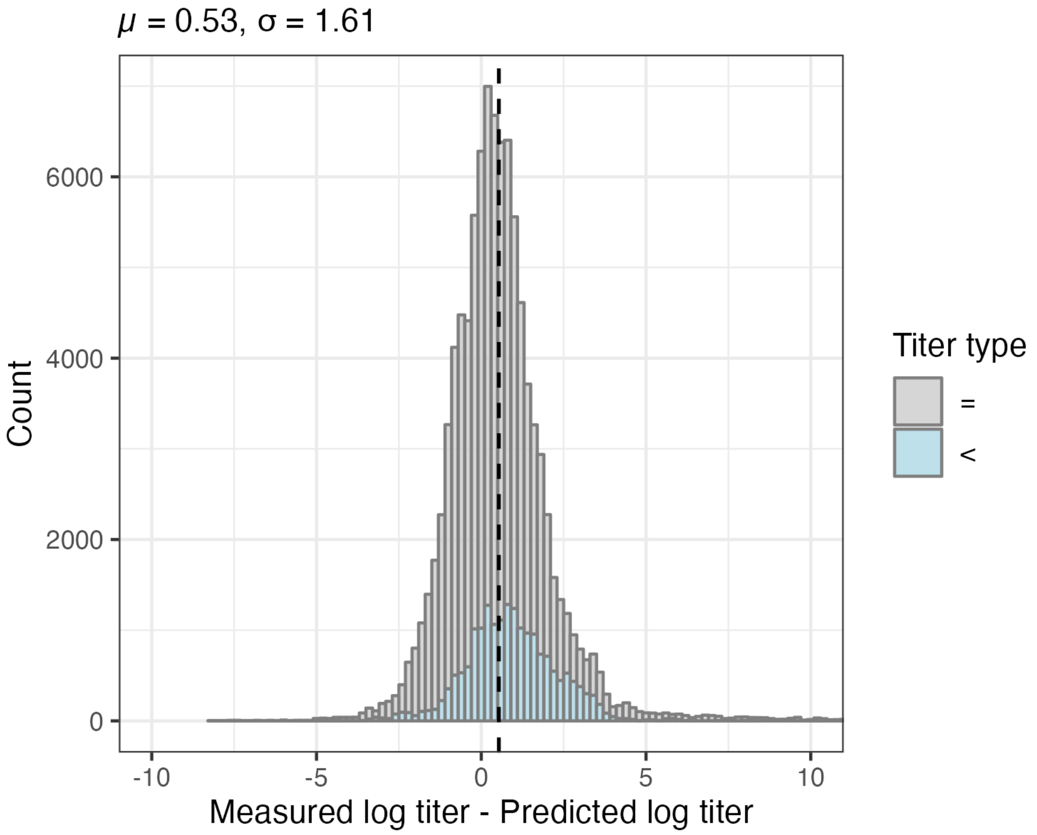


**Figure S20: Histogram of differences between measured and cross-validation predicted $\log_2$ titers.** For non-detectable measured titers, differences between predicted and measured titers were imputed as described in the methods section, and are indicated in light blue. Differences for detectable measured titers are indicated in gray. Predicted titers are on average 0.53 lower than measured titers with a standard deviation of 1.61 on the $\log_2$ scale. The black dashed line indicates the mean.

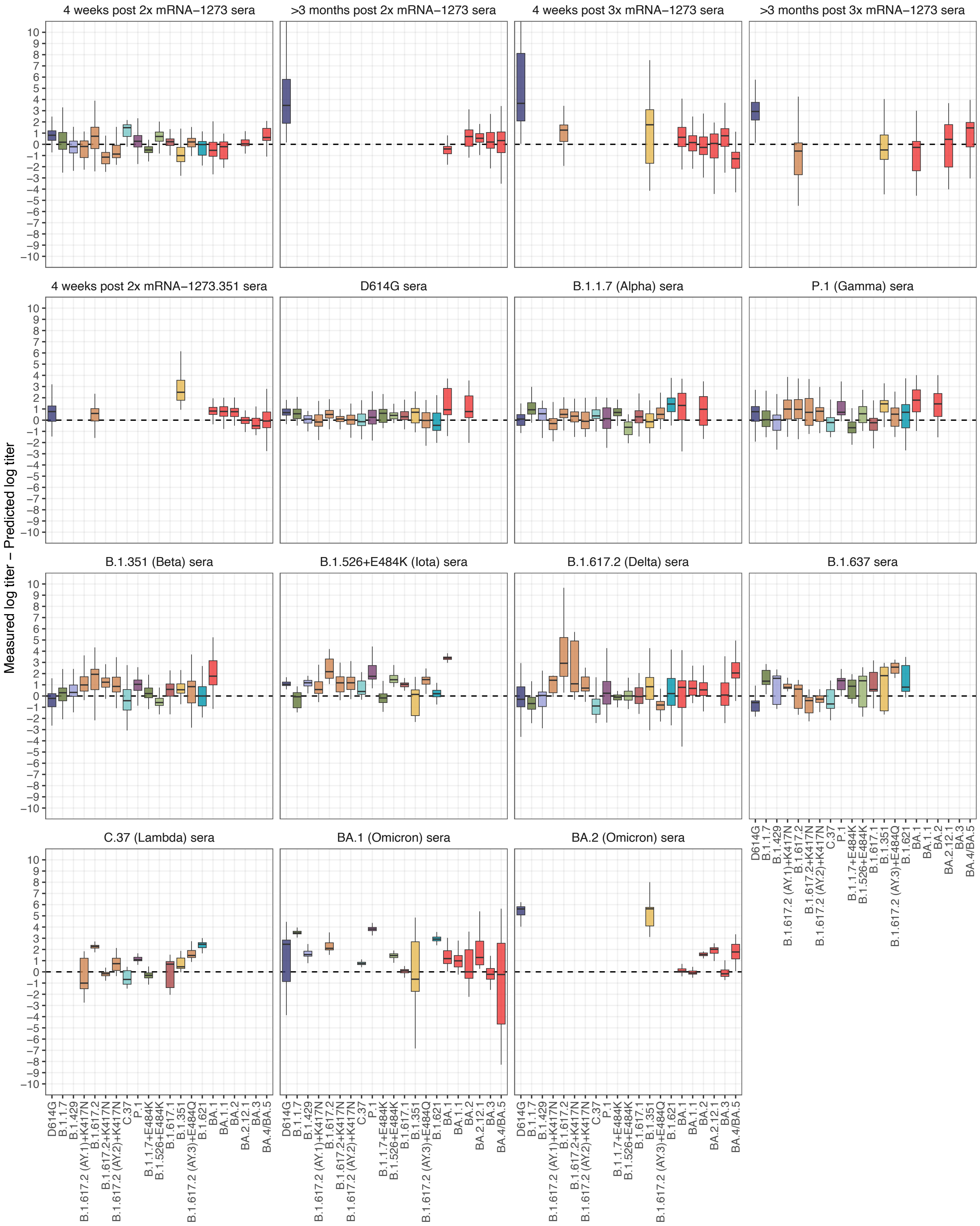


**Figure S21: Measured minus cross-validation predicted $log_2$ titers split by serum group and variant.** The difference between predicted and measured $log_2$ titers is shown split by variant and serum group. The box plot indicates the median and 25th and 75th percentile.

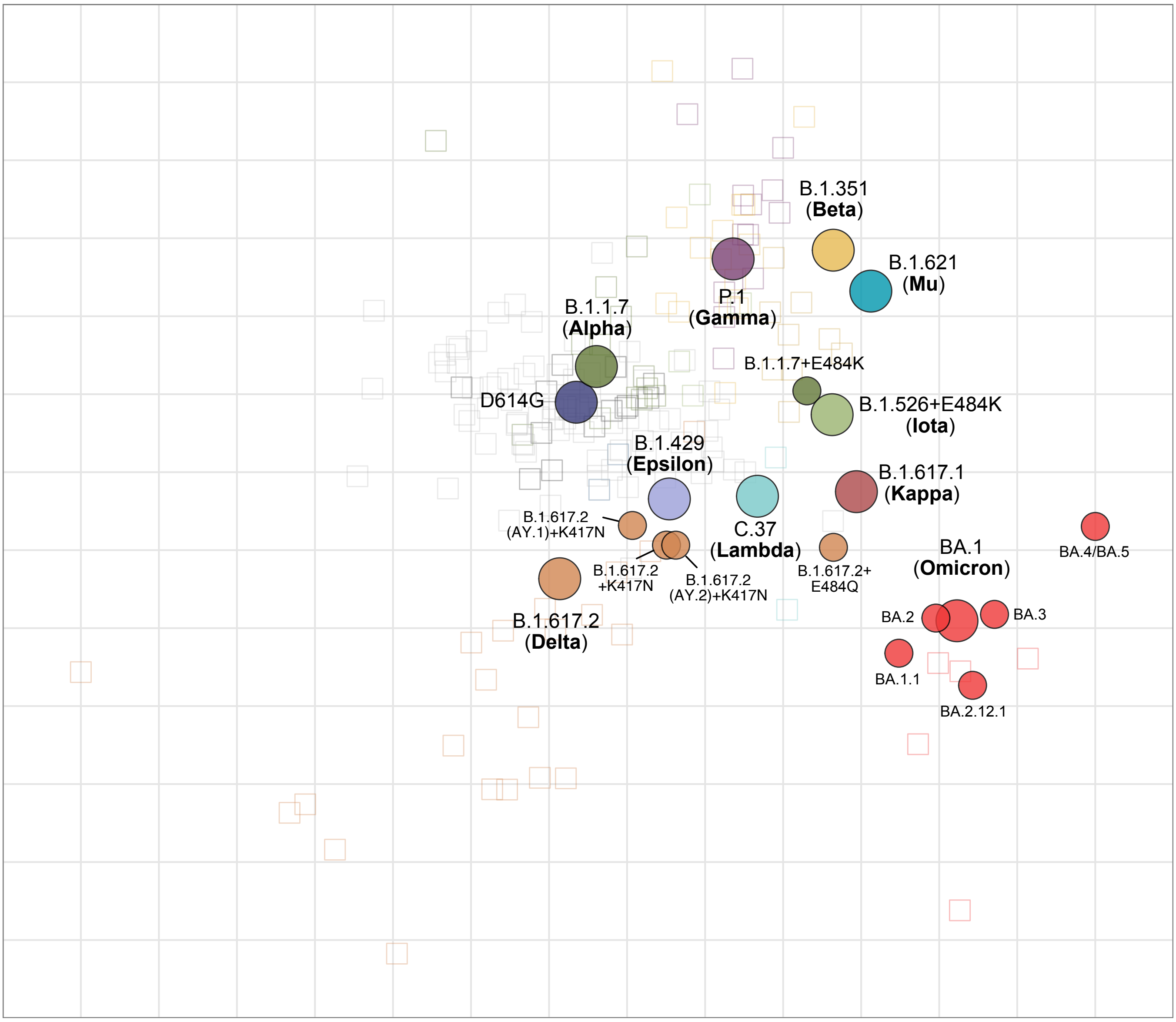


**Figure S22: Zoomed out version of the antigenic map shown in Fig. 3.** Variants are shown as circles, sera as squares, sera colors correspond to the color of the eliciting variant. Variants with additional substitutions from a root variant are denoted by smaller circles, in the color of their root variant. The x and y-axes both represent antigenic distance, with one grid square corresponding to one two-fold serum dilution in the neutralization assay.

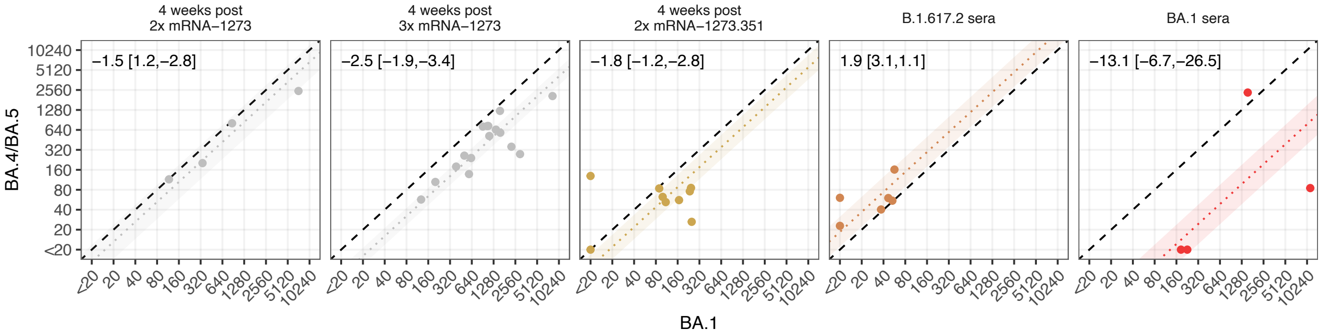


**Figure S23: Scatterplots comparing serum titers against variants BA.1 and BA.4/BA.5.** The dotted line shows a line with slope 1 and intercept equal to the fold-change point estimate of difference between the variants on the x and y-axes; shaded regions represent the 95% HDI for these estimates. The dashed black line shows the line of equality. Numbers in the top left corner show the associated point estimates and HDI expressed as fold-changes.

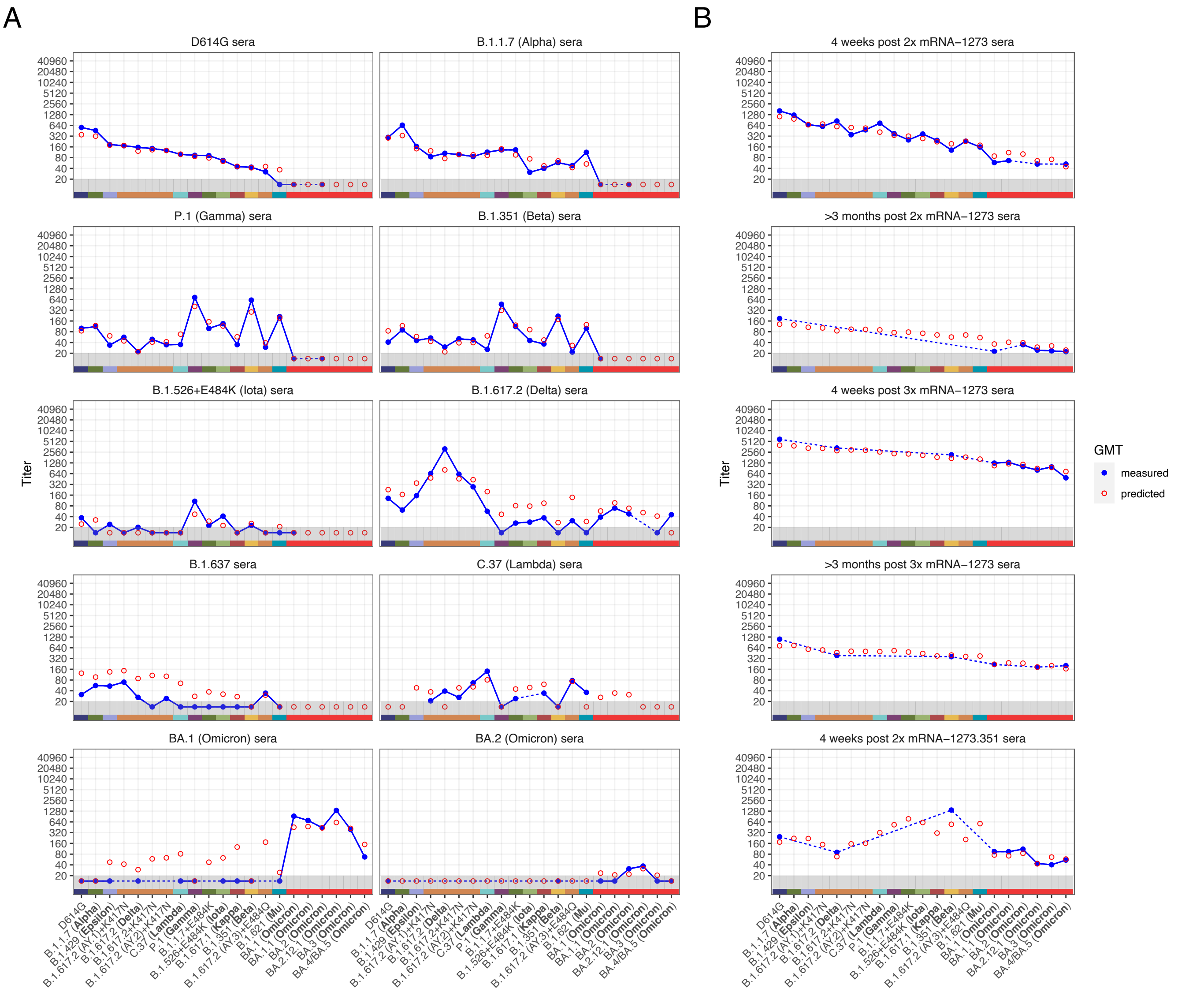


**Figure S24: Measured vs antibody landscapes-predicted GMTs for each serum group**. Blue points and the line show the measured GMT for each variant, after accounting for individual effects, as described above and plotted also in Fig. 1, which would otherwise bias the GMT for variants not titrated against all sera. Red outlined circles show GMT predictions according to the map and antibody landscape fits as described in Materials and Methods, ‘Construction of the antibody landscapes’. Points in the gray region at the bottom of the plot show where the measured or predicted GMT has fallen below the assay detection threshold of 20.

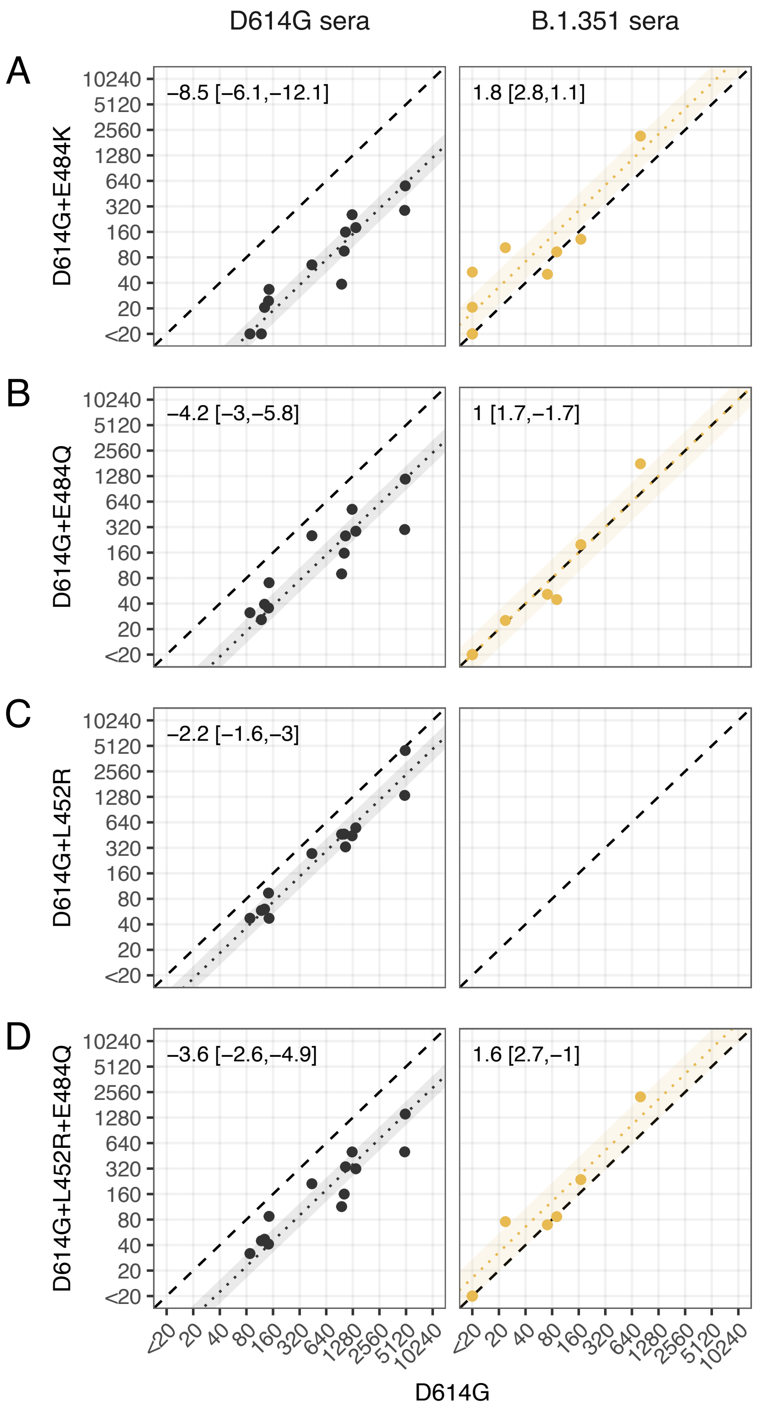


**Figure S25: Scatterplots showing the comparison of titers of the single and double mutants D614G+E484K, D614G+E484Q, D614G+L452R, and D614G+L452R+E484Q to titers of the root virus D614G for the D614G and B.1.351 sera.** The dotted line shows a line with slope 1 and intercept equal to the fold-change point estimate of difference between the variants on the x and y-axes; shaded regions represent the 95% HDI for these estimates. The dashed black line shows the line of equality. Numbers in the top left corner show the associated point estimates and HDI expressed as fold-changes.

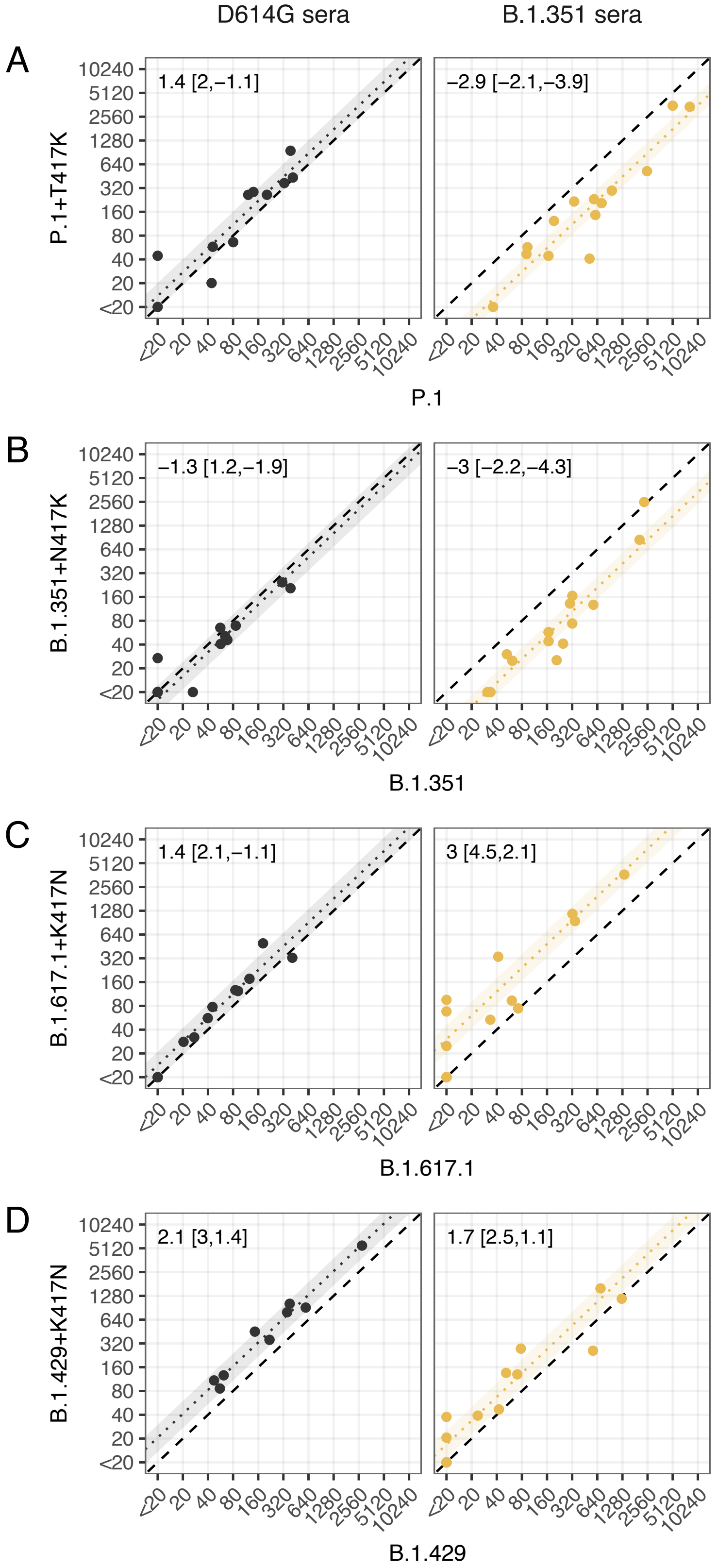


**Figure S26: Scatterplots showing the effect of introducing substitutions at 417 into different variant backgrounds.** The dotted line shows a line with slope 1 and intercept equal to the fold-change point estimate of difference between the variants on the x and y-axes; shaded regions represent the 95% HDI for these estimates. The dashed black line shows the line of equality. Numbers in the top left corner show the associated point estimates and HDI expressed as fold-changes.

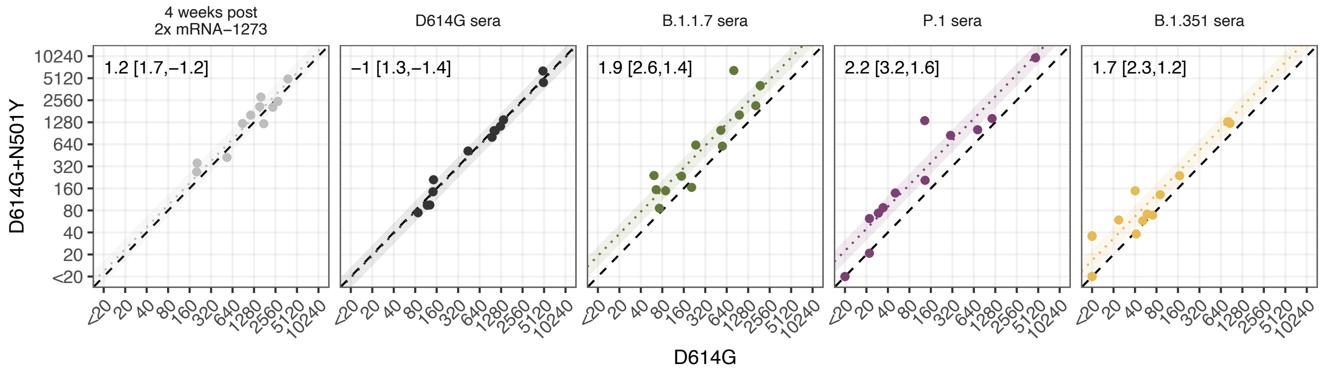


**Figure S27: Scatterplots showing the comparison of titers of the single mutants D614G+N501Y to titers of the root virus D614G.** The dotted line shows a line with slope 1 and intercept equal to the fold-change point estimate of difference between the variants on the x and y-axes; shaded regions represent the 95% HDI for these estimates. The dashed black line shows the line of equality. Numbers in the top left corner show the associated point estimates and HDI expressed as fold-changes.

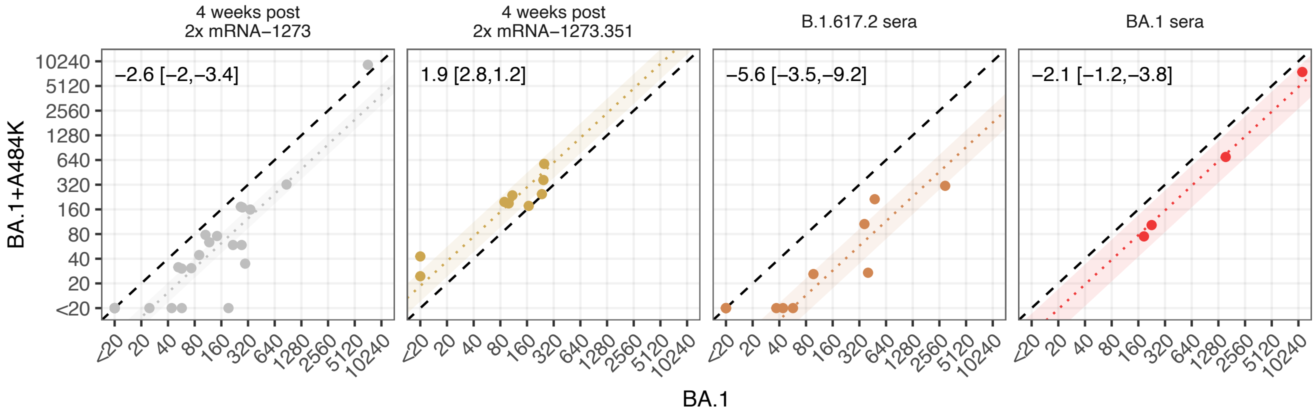


**Figure S28: Scatterplots showing the effect of introducing substitutions at A484K into the BA.1 variant background.** The dotted line shows a line with slope 1 and intercept equal to the fold-change point estimate of difference between the variants on the x and y-axes; shaded regions represent the 95% HDI for these estimates. The dashed black line shows the line of equality. Numbers in the top left corner show the associated point estimates and HDI expressed as fold-changes.

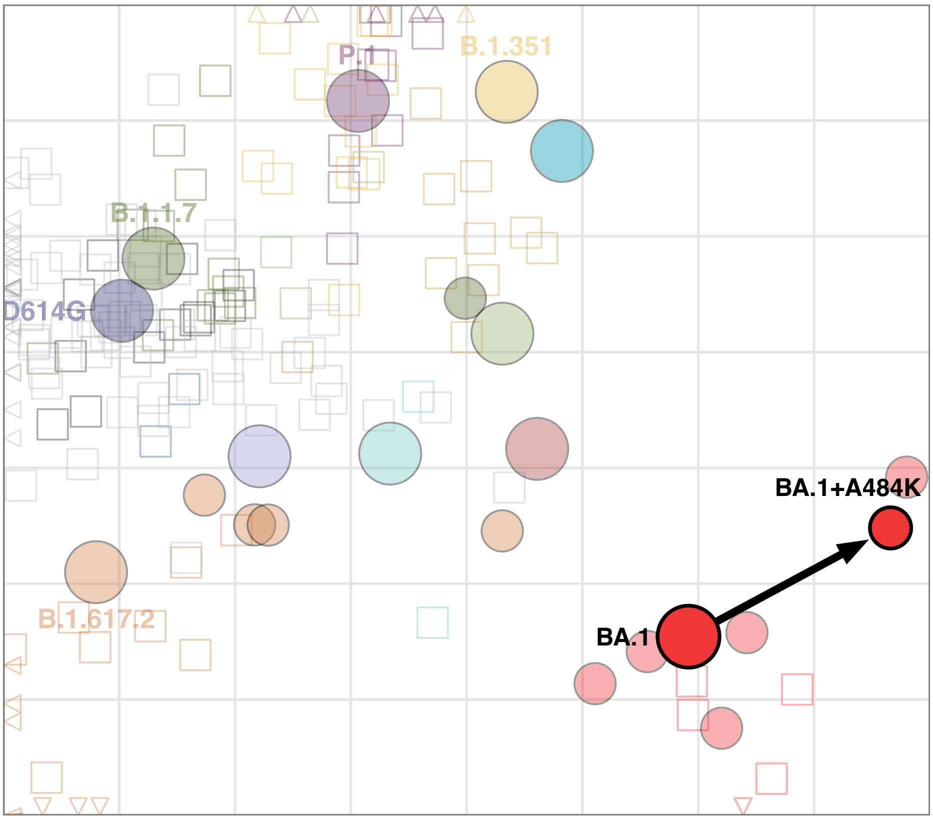


**Figure S29: Two-dimensional antigenic map including the BA.1+A484K mutant.** The antigenic map with the laboratory made BA.1+A484K mutant included. This is made from the same data as the antigenic map in Fig. 5C but optimized in two dimensions. The arrow points from the antigenic position of the root virus to that of the laboratory-generated variant.

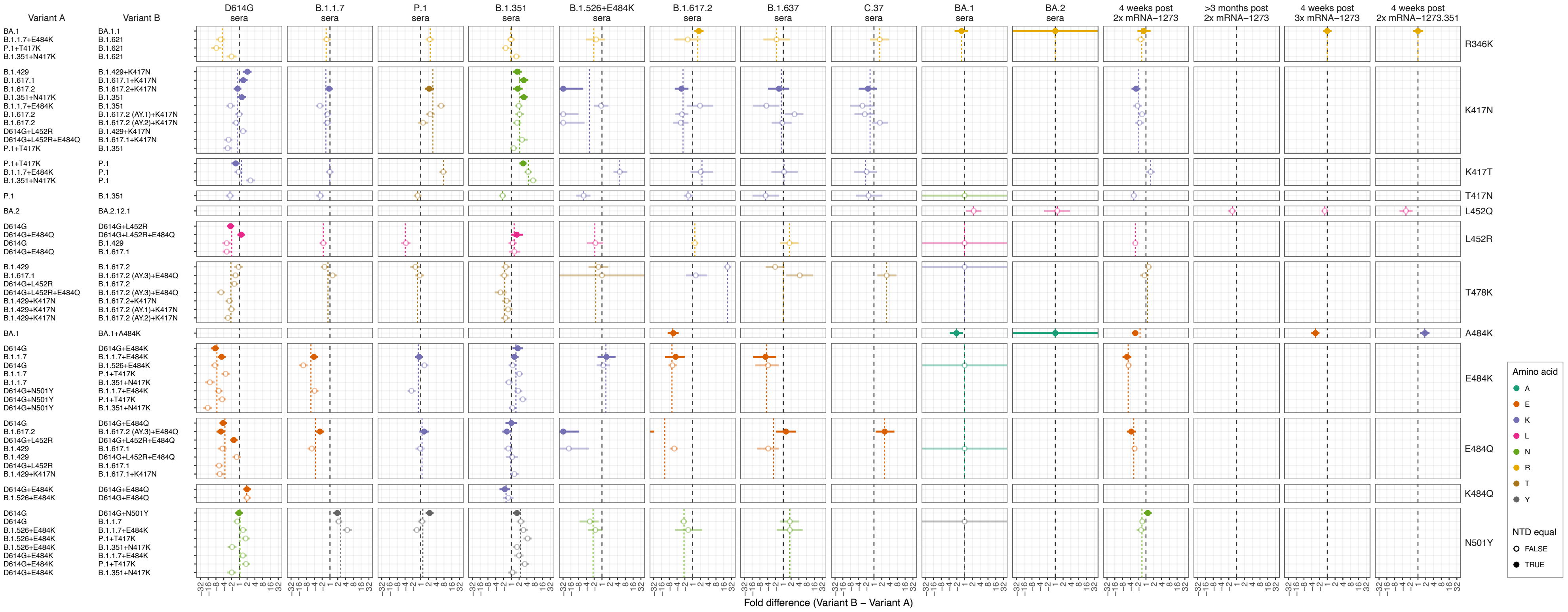


**Figure S30: Effect of pairwise amino acid differences on reactivity to different serum groups.** This plot compares the average fold difference in titer between different pairs of variants that differ by only a single amino acid substitution in the RBD, and includes all pairwise examples and serum groups, even where less data is available. Comparisons are grouped by serum group (panel columns) and corresponding RBD difference (panel rows). In each panel, the circle represents the estimate for the average fold difference in titer between variant A and variant B, as defined on the left-hand side of the plot, while lines extend to indicate the 95% highest density interval for this estimate. The black dashed line marks a fold difference in titer of 1 (no difference), while the colored dashed line indicates the mean fold difference between all pairs of variants with the same substitution in the RBD. Points and lines are colored according to the corresponding amino acid in the variant homologous to the serum group, at the position in the RBD where the pair of variants compared differ. Filled circles indicate where pairs of variants have no additional amino acid differences in the NTD, often because one was generated as an artificial mutant. In contrast, open circles indicate pairs of variants with amino acid differences in the NTD, in addition to the RBD amino acid difference listed. Details of how fold-difference estimates and highest density intervals were calculated are described in Materials and Methods.

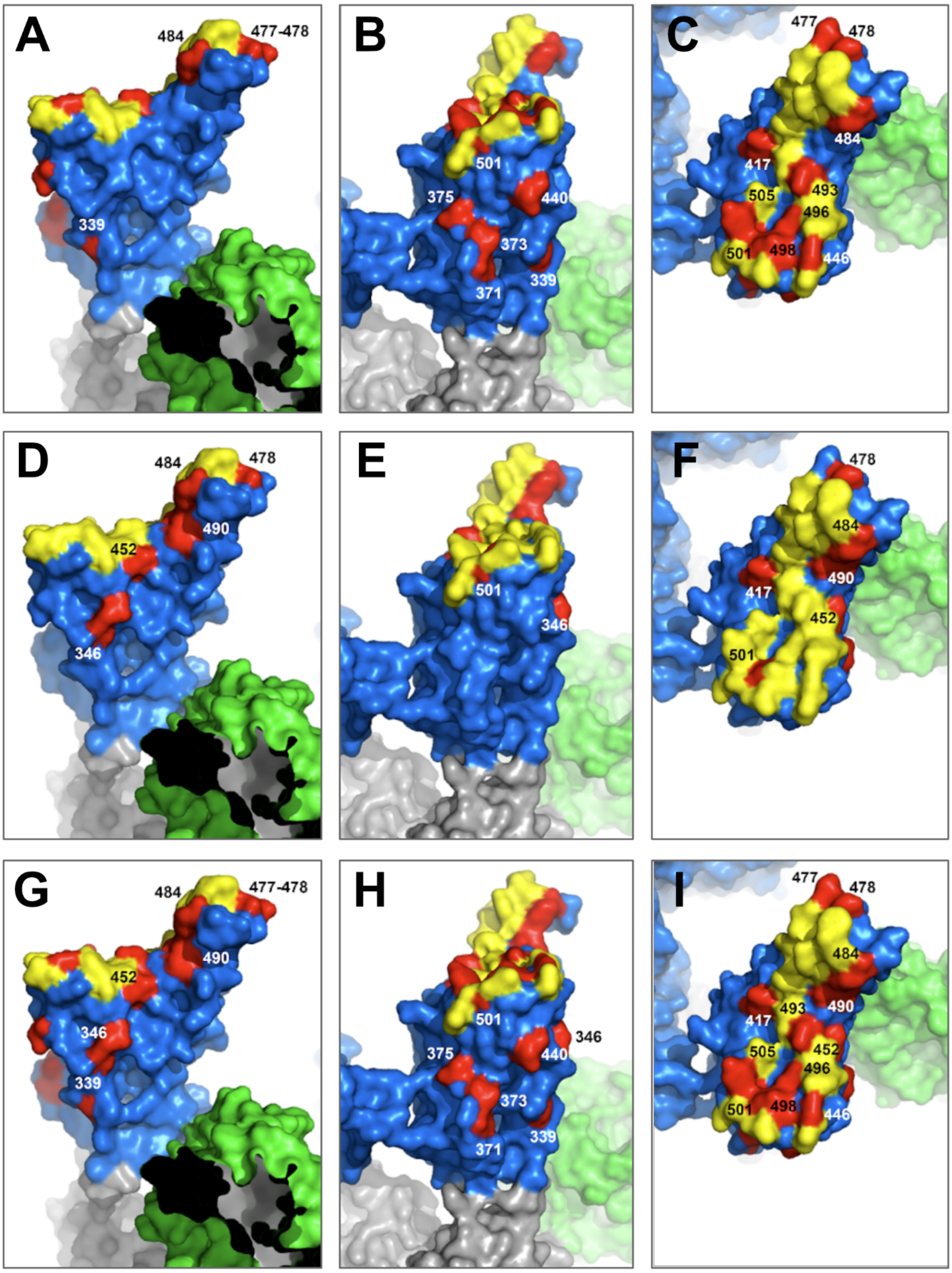


**Figure S31: Visualization of RBD substitutions present in variants in this study on the SARS-CoV-2 spike structure.** RBD in blue, RBD-ACE2 interface residues in yellow, NTD in green, variant substitutions in red. Substitutions labeled only on the "up" conformation of the RBD. A-C) Positions with substitutions in the BA.1 variant. D-F) Substitutions of the positions with substitutions in non-Omicron variants from this study. G-I) Positions with substitutions in all variants in this study. A, D, G) Solvent-accessible front of the RBD. B, E, H) Side of the RBD (at the RBD-RBD interface when the RBD is in the "closed" conformation). C, F, I) RBD-ACE2 interaction interface of the RBD. The structure was produced using Modeller, by combining data from three structures with protein data bank accessions 7KNB, 7C2L, 7L2C.

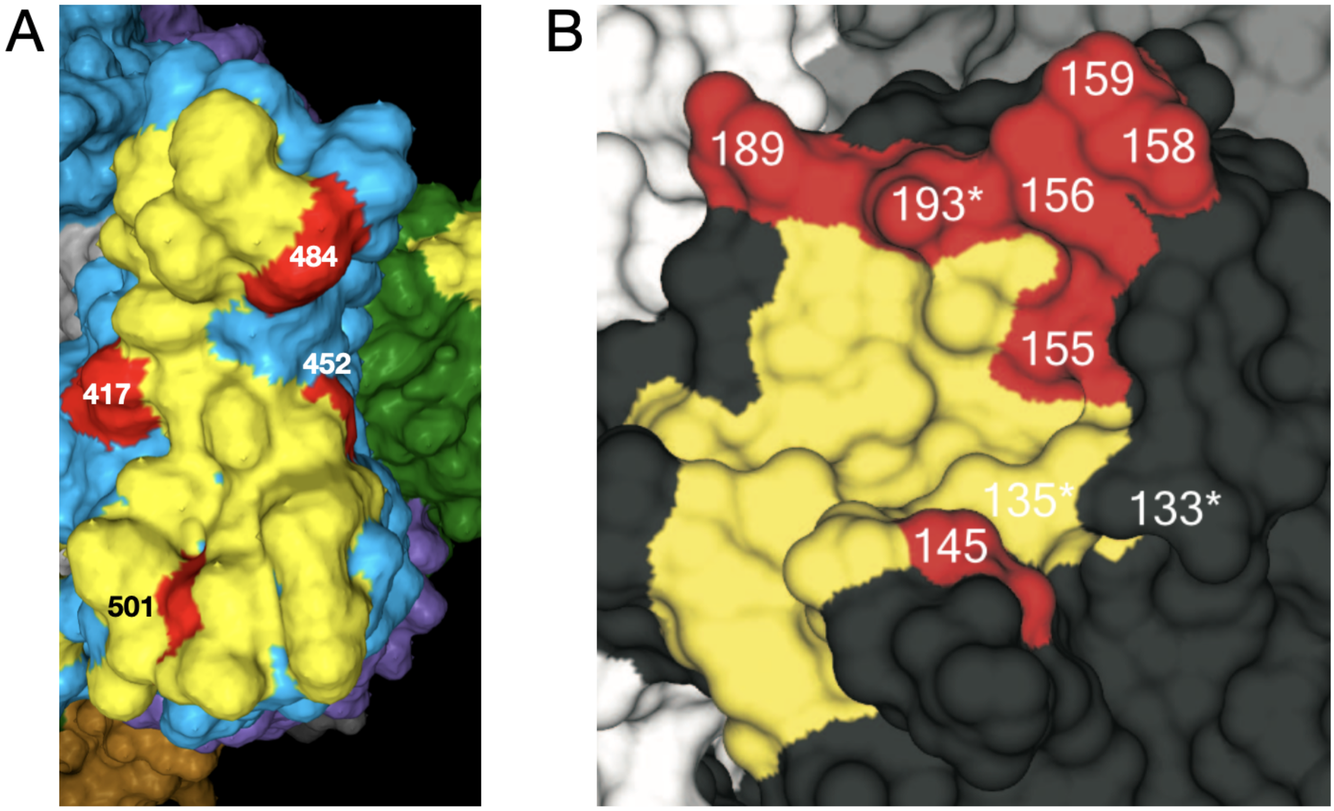


**Figure S32: Comparison of putative antigenic substitutions in SARS-CoV-2 and Influenza A/H3N2.** A) SARS-CoV-2 RBD with key positions 417, 452, 484, and 501 marked in red, ACE2 interaction interface is shown in yellow. B) Influenza A/H3N2 receptor binding site, in yellow, with key antigenic positions marked in red (133, 135, 145, 155, 156, 158, 159, 189, and 193). Figure taken from Koel et al., 2013 (*40*)

| Sample Group | Agency / Cohort[#] | n | Lead Contacts | Collection Location | Infection Stage at Collection[$] | First infection criteria |
|---|---|---|---|---|---|---|
| 4 weeks post 2x mRNA-1273 | Moderna / phase 1 study<br>Moderna / COVE phase 3 trial | 28<br>41* | Pajon | United States | Non infected | - |
| >3 months post 2x mRNA-1273 | NIH / DMID 21-0012 | 16 | Beigel | United States | Non infected | - |
| 4 weeks post 3x mRNA-1273 | NIH / DMID 21-0012 | 26 | Beigel | United States | Non infected | - |
| 6 months post 3x mRNA-1273 | NIH / CoVAIL | 8 | Beigel | United States | Non infected | - |
| 4 weeks post mRNA-1273.351 | NIH / DMID 21-0002 | 11 | Beigel | United States | Non infected | - |
| D614G sera | CoVPN | 15 | Hural | United States | Convalescent | 1. |
| B.1.1.7 sera | BC CDC | 14 | Jassem | Canada | Convalescent | 1. |
| B.1.351 sera | CoVPN | 19 | Hural | Africa | Convalescent | 1. |
| P.1 sera | BC CDC | 16 | Jassem | Canada | Convalescent | 1. |
| | UW-Madison | 1 | Kawaoka | Japan | Active infection | 1. |
| B.1.617.2 sera | BC CDC | 17 | Jassem | Canada | Convalescent | 1. |
| | UW-Madison | 2 | Kawaoka | Japan | Active infection | 1. |
| | CoVPN | 9 | Hural | United States | Convalescent | 1. |
| B.1.526+E484K sera | CDC / C-HEaRT | 2 | Dawood & Veguilla | United States | Convalescent | 2. |
| | Icahn School of Medicine at Mount Sinai | 4 | Krammer & Simon | United States | Convalescent/Longevity | 1. |
| C.37 sera | St. Jude Children's Research Hospital | 4 | Webby | Peru | Convalescent | 3. |
| B.1.637 sera | CDC / C-HEaRT | 3 | Dawood & Veguilla | United States | Convalescent | 2. |
| BA.1 sera | Charité / VariPath | 3 | Corman & Jeworowski | Germany | Convalescent | 1. |
| | BC CDC | 4 | Jassem | Canada | Convalescent | 1. |
| BA.2 sera | Charité / VariPath | 1 | Corman & Jeworowski | Germany | Convalescent | 1. |

**Table S1: Description of sera used in this study.** [#]BC CDC-BC Centre for Disease Control; CoVPN- COVID-19 Prevention Network; UW-Madison- University of Wisconsin, Madison; CDC-Centers for Disease Control and Prevention; C-HEaRT- Coronavirus Household Evaluation and Respiratory Testing Cohort; Charité: Charité – Universitätsmedizin Berlin, Institute of Virology. [$]Defined by days since symptom onset or first positive diagnosis. Active infection: <20 days; Convalescent: ≥28 days and <60 days; Longevity: ≥60 days. *Note that only four of these samples contribute to the general analyses, with others contributing to specific mutant sub-analyses as described in “Specimens and study cohorts”. First infection criteria: 1. No previous vaccination or infection self-reported or in patient records. 2. Participants submitted weekly mid-turbinate nasal swabs regardless of symptoms throughout the study period. Swabs were tested for SARS-CoV-2 by RT-PCR and positive samples were subsequently sequenced. Based on US surveillance data, neither variant (B.1.525+E484K and B.1.637) circulated substantially before the study period. 3. Early samples after symptom onset were NP ELISA negative, later samples were subsequently sequence confirmed C.37 infections.

| | 5 | 13 | NTD 18 | NTD 19 | NTD 20 | NTD 24 | NTD 25 | NTD 26 | NTD 27 | NTD 67 | NTD 69 | NTD 70 | NTD 75 | NTD 76 | NTD 80 | NTD 95 | NTD 138 | NTD 142 | NTD 143 | NTD 143.1 | NTD 144 | NTD 145 | NTD 152 | NTD 154 | NTD 156 | NTD 157 |
|---|---|---|---|---|---|---|---|---|---|---|---|---|---|---|---|---|---|---|---|---|---|---|---|---|---|---|
| D614G | L | S | L | T | T | L | P | P | A | A | H | V | G | T | D | T | D | G | V | - | Y | Y | W | E | E | F |
| B.1.1.7 | . | . | . | . | . | . | . | . | . | . | - | - | . | . | . | . | . | . | . | - | - | . | . | . | . | . |
| B.1.1.7+E484K | . | . | . | . | . | . | . | . | . | . | - | - | . | . | . | . | . | . | . | - | - | . | . | . | . | . |
| P.1 | . | . | F | . | N | . | . | S | . | . | . | . | . | . | . | . | Y | . | . | - | . | . | . | . | . | . |
| B.1.351 | . | . | F | . | . | . | . | . | . | . | . | . | . | . | A | . | . | . | . | - | . | . | . | . | . | . |
| B.1.429 | . | I | . | . | . | . | . | . | . | . | . | . | . | . | . | . | . | . | . | - | . | . | C | . | . | . |
| B.1.526+E484K | F | . | . | . | . | . | . | . | . | . | . | . | . | . | . | I | . | . | . | - | . | . | . | . | . | . |
| B.1.617.1 | . | . | . | . | . | . | . | . | . | . | . | . | . | . | . | . | . | D | . | - | . | . | . | K | . | . |
| B.1.617.2 | . | . | . | R | . | . | . | . | . | . | . | . | . | . | . | . | . | D | . | - | . | . | . | . | - | - |
| B.1.617.2+K417N | . | . | . | R | . | . | . | . | . | . | . | . | . | . | . | . | . | D | . | - | . | . | . | . | - | - |
| B.1.617.2 (AY.1)+K417N | . | . | . | R | . | . | . | . | . | . | . | . | . | . | . | I | . | D | . | - | . | . | . | . | - | - |
| B.1.617.2 (AY.2)+K417N | . | . | . | R | . | . | . | . | . | . | . | F | . | . | . | . | . | D | . | - | . | . | . | . | - | - |
| B.1.617.2 (AY.3)+E484Q | . | . | . | R | . | . | . | . | . | . | . | . | . | . | . | . | . | D | . | - | . | . | . | . | - | - |
| B.1.621 | . | . | . | . | . | . | . | . | . | . | . | . | . | . | . | I | . | . | V | T | S | N | . | . | . | . |
| C.37 | . | . | . | . | . | . | . | . | . | . | . | . | V | I | . | . | . | . | . | - | . | . | . | . | . | . |
| BA.1 | . | . | . | . | . | . | . | . | . | V | - | - | . | . | . | I | . | D | - | - | - | - | . | . | . | . |
| BA.1.1 | . | . | . | . | . | . | . | . | . | V | - | - | . | . | . | I | . | D | - | - | - | - | . | . | . | . |
| BA.2 | . | . | . | I | . | - | - | - | S | . | . | . | . | . | . | . | . | D | . | - | . | . | . | . | . | . |
| BA.2.12.1 | . | . | . | I | . | S | - | - | - | . | . | . | . | . | . | . | . | D | . | - | . | . | . | . | . | . |
| BA.3 | . | . | . | . | . | . | . | . | . | V | - | - | . | . | . | I | . | D | - | - | - | - | . | . | . | . |
| BA.4/BA.5 | . | . | . | I | . | S | - | - | - | . | . | . | . | . | . | . | . | D | . | - | . | . | . | . | . | . |

| | NTD 158 | NTD 190 | NTD 211 | NTD 212 | NTD 213 | NTD 214 | NTD 214.1 | NTD 214.2 | NTD 214.3 | NTD 215 | NTD 222 | NTD 241 | NTD 242 | NTD 243 | NTD 246 | NTD 247 | NTD 248 | NTD 249 | NTD 250 | NTD 251 | NTD 252 | NTD 253 | NTD 258 |
|---|---|---|---|---|---|---|---|---|---|---|---|---|---|---|---|---|---|---|---|---|---|---|---|
| D614G | R | R | N | L | V | R | - | - | - | D | A | L | L | A | R | S | Y | L | T | P | G | D | W |
| B.1.1.7 | . | . | . | . | . | . | - | - | - | . | . | . | . | . | . | . | . | . | . | . | . | . | . |
| B.1.1.7+E484K | . | . | . | . | . | . | - | - | - | . | . | . | . | . | . | . | . | . | . | . | . | . | . |
| P.1 | . | S | . | . | . | . | - | - | - | . | . | . | . | . | . | . | . | . | . | . | . | . | . |
| B.1.351 | . | . | . | . | . | . | - | - | - | G | . | - | - | - | I | . | . | . | . | . | . | . | . |
| B.1.429 | . | . | . | . | . | . | - | - | - | . | . | . | . | . | . | . | . | . | . | . | . | . | . |
| B.1.526+E484K | . | . | . | . | . | . | - | - | - | . | . | . | . | . | . | . | . | . | . | . | . | G | . |
| B.1.617.1 | . | . | . | . | . | . | - | - | - | . | . | . | . | . | . | . | . | . | . | . | . | . | . |
| B.1.617.2 | G | . | . | . | . | . | - | - | - | . | . | . | . | . | . | . | . | . | . | . | . | . | . |
| B.1.617.2+K417N | G | . | . | . | . | . | - | - | - | . | . | . | . | . | . | . | . | . | . | . | . | . | . |
| B.1.617.2 (AY.1)+K417N | G | . | . | . | . | . | - | - | - | . | . | . | . | . | . | . | . | . | . | . | . | . | L |
| B.1.617.2 (AY.2)+K417N | G | . | . | . | . | . | - | - | - | . | V | . | . | . | . | . | . | . | . | . | . | . | . |
| B.1.617.2 (AY.3)+E484Q | G | . | . | . | . | . | - | - | - | . | . | . | . | . | . | . | . | . | . | . | . | . | . |
| B.1.621 | . | . | . | . | . | . | - | - | - | . | . | . | . | . | . | . | . | . | . | . | . | . | . |
| C.37 | . | . | . | . | . | . | - | - | - | . | . | . | . | . | N | - | - | - | - | - | - | - | . |
| BA.1 | . | . | - | I | . | R | E | P | E | . | . | . | . | . | . | . | . | . | . | . | . | . | . |
| BA.1.1 | . | . | - | I | . | R | E | P | E | . | . | . | . | . | . | . | . | . | . | . | . | . | . |
| BA.2 | . | . | . | . | G | . | - | - | - | . | . | . | . | . | . | . | . | . | . | . | . | . | . |
| BA.2.12.1 | . | . | . | . | G | . | - | - | - | . | . | . | . | . | . | . | . | . | . | . | . | . | . |
| BA.3 | . | . | - | I | . | . | - | - | - | . | . | . | . | . | . | . | . | . | . | . | . | . | . |
| BA.4/BA.5 | . | . | . | . | G | . | - | - | - | . | . | . | . | . | . | . | . | . | . | . | . | . | . |

| | RBD 339 | RBD 346 | RBD 371 | RBD 373 | RBD 375 | RBD 376 | RBD 405 | RBD 408 | RBD 417 | RBD 440 | RBD 446 | RBD 452 | RBD 477 | RBD 478 | RBD 484 | RBD 486 | RBD 490 | RBD 493 | RBD 496 | RBD 498 | RBD 501 | RBD 505 |
|---|---|---|---|---|---|---|---|---|---|---|---|---|---|---|---|---|---|---|---|---|---|---|
| D614G | G | R | S | S | S | T | D | R | K | N | G | L | S | T | E | F | F | Q | G | Q | N | Y |
| B.1.1.7 | . | . | . | . | . | . | . | . | . | . | . | . | . | . | . | . | . | . | . | . | Y | . |
| B.1.1.7+E484K | . | . | . | . | . | . | . | . | . | . | . | . | . | . | K | . | . | . | . | . | Y | . |
| P.1 | . | . | . | . | . | . | . | . | T | . | . | . | . | . | K | . | . | . | . | . | Y | . |
| B.1.351 | . | . | . | . | . | . | . | . | N | . | . | . | . | . | K | . | . | . | . | . | Y | . |
| B.1.429 | . | . | . | . | . | . | . | . | . | . | . | R | . | . | . | . | . | . | . | . | . | . |
| B.1.526+E484K | . | . | . | . | . | . | . | . | . | . | . | . | . | . | K | . | . | . | . | . | . | . |
| B.1.617.1 | . | . | . | . | . | . | . | . | . | . | . | R | . | . | Q | . | . | . | . | . | . | . |
| B.1.617.2 | . | . | . | . | . | . | . | . | . | . | . | R | . | K | . | . | . | . | . | . | . | . |
| B.1.617.2+K417N | . | . | . | . | . | . | . | . | N | . | . | R | . | K | . | . | . | . | . | . | . | . |
| B.1.617.2 (AY.1)+K417N | . | . | . | . | . | . | . | . | N | . | . | R | . | K | . | . | . | . | . | . | . | . |
| B.1.617.2 (AY.2)+K417N | . | . | . | . | . | . | . | . | N | . | . | R | . | K | . | . | . | . | . | . | . | . |
| B.1.617.2 (AY.3)+E484Q | . | . | . | . | . | . | . | . | . | . | . | R | . | K | Q | . | . | . | . | . | . | . |
| B.1.621 | . | K | . | . | . | . | . | . | . | . | . | . | . | . | K | . | . | . | . | . | Y | . |
| C.37 | . | . | . | . | . | . | . | . | . | . | . | Q | . | . | . | . | S | . | . | . | . | . |
| BA.1 | D | . | L | P | F | . | . | . | N | K | S | . | N | K | A | . | . | R | S | R | Y | H |
| BA.1.1 | D | K | L | P | F | . | . | . | N | K | S | . | N | K | A | . | . | R | S | R | Y | H |
| BA.2 | D | . | F | P | F | A | N | S | N | K | . | . | N | K | A | . | . | R | . | R | Y | H |
| BA.2.12.1 | D | . | F | P | F | A | N | S | N | K | . | Q | N | K | A | . | . | R | . | R | Y | H |
| BA.3 | D | . | F | P | F | . | N | . | N | K | S | . | N | K | A | . | . | R | . | R | Y | H |
| BA.4/BA.5 | D | . | F | P | F | A | N | S | N | K | . | R | N | K | A | V | . | . | . | R | Y | H |

| | 547 | 570 | 655 | 679 | 681 | 701 | 704 | 716 | 764 | 796 | 856 | 859 | 950 | 954 | 969 | 981 | 982 | 1027 | 1071 | 1118 |
|---|---|---|---|---|---|---|---|---|---|---|---|---|---|---|---|---|---|---|---|---|
| D614G | T | A | H | N | P | A | S | T | N | D | N | T | D | Q | N | L | S | T | Q | D |
| B.1.1.7 | . | D | . | . | H | . | . | I | . | . | . | . | . | . | . | . | A | . | . | H |
| B.1.1.7+E484K | . | D | . | . | H | . | . | I | . | . | . | . | . | . | . | . | A | . | . | H |
| P.1 | . | . | Y | . | . | . | . | . | . | . | . | . | . | . | . | . | . | I | . | . |
| B.1.351 | . | . | . | . | . | V | . | . | . | . | . | . | . | . | . | . | . | . | . | . |
| B.1.429 | . | . | . | . | . | . | . | . | . | . | . | . | . | . | . | . | . | . | . | . |
| B.1.526+E484K | . | . | . | . | . | V | . | . | . | . | . | . | . | . | . | . | . | . | . | . |
| B.1.617.1 | . | . | . | . | R | . | . | . | . | . | . | . | . | . | . | . | . | . | H | . |
| B.1.617.2 | . | . | . | . | R | . | . | . | . | . | . | . | N | . | . | . | . | . | . | . |
| B.1.617.2+K417N | . | . | . | . | R | . | . | . | . | . | . | . | N | . | . | . | . | . | . | . |
| B.1.617.2 (AY.1)+K417N | . | . | . | . | R | . | . | . | . | . | . | . | N | . | . | . | . | . | . | . |
| B.1.617.2 (AY.2)+K417N | . | . | . | . | R | . | . | . | . | . | . | . | N | . | . | . | . | . | . | . |
| B.1.617.2 (AY.3)+E484Q | . | . | . | . | R | . | . | . | . | . | . | . | N | . | . | . | . | . | . | . |
| B.1.621 | . | . | . | . | H | . | . | . | . | . | . | . | N | . | . | . | . | . | . | . |
| C.37 | . | . | . | . | . | . | . | . | . | . | . | N | . | . | . | . | . | . | . | . |
| BA.1 | K | . | Y | K | H | . | . | . | K | Y | K | . | . | H | K | F | . | . | . | . |
| BA.1.1 | K | . | Y | K | H | . | . | . | K | Y | K | . | . | H | K | F | . | . | . | . |
| BA.2 | . | . | Y | K | H | . | . | . | K | Y | . | . | . | H | K | . | . | . | . | . |
| BA.2.12.1 | . | . | Y | K | H | . | L | . | K | Y | . | . | . | H | K | . | . | . | . | . |
| BA.3 | . | . | Y | K | H | . | . | . | K | Y | . | . | . | H | K | . | . | . | . | . |
| BA.4/BA.5 | . | . | Y | K | H | . | . | . | K | Y | . | . | . | H | K | . | . | . | . | . |

**Table S2: A comparison of sequence differences of variants used in this study.** This table displays the amino acid differences compared to D614G in the spike protein for each of the variants. The coloring is according to the Lesk scheme which categorizes amino acids based on physico-chemical similarity (*69*) as follows; orange: small nonpolar, green: hydrophobic, magenta: polar, red: negatively charged, blue: positively charged.
